# Isoleucine binding and regulation of *Escherichia coli* and *Staphylococcus aureus* threonine dehydratase (IlvA)

**DOI:** 10.1101/2025.03.06.641827

**Authors:** Mi-Kyung Yun, Chitra Subramanian, Karen Miller, Pamela Jackson, Christopher D. Radka, Charles O. Rock

## Abstract

In *Staphylococcus aureus*, the branched-chain amino acid biosynthetic pathway provides essential intermediates for membrane biosynthesis. Threonine deaminase (IlvA) is the first enzyme in the pathway, and isoleucine feedback-regulates the enzyme in *Escherichia coli*. These studies on *E. coli* IlvA (EcIlvA) introduced the concept of allosteric regulation. To investigate the regulation of *S. aureus* IlvA (SaIlvA), we first conducted additional studies on EcIlvA. The previously determined crystal structure of EcIlvA revealed a tetrameric assembly of protomers, each with catalytic and regulatory domains, but the structural basis of isoleucine regulation was not characterized. Here, we present the crystal structure of the EcIlvA regulatory domain bound to isoleucine, which reveals the isoleucine binding site and conformational changes that initiate at Phe352 and propagate 23 Å across the domain. This suggests an allosteric pathway that extends to the active site of the adjacent protomer, mediating regulation across the protomer-protomer interface. The EcIlvA(F352A) mutant binds isoleucine but is feedback-resistant due to the absence of the initiating Phe352. In contrast, SaIlvA is not feedback-regulated by isoleucine and does not bind it. The structure of the SaIlvA regulatory domain reveals a different organization that lacks the isoleucine binding site. Other potential allosteric inhibitors of SaIlvA, including phospholipid intermediates, do not affect enzyme activity. We propose that the absence of feedback inhibition in SaIlvA is due to its role in membrane biosynthesis. These findings enhance our understanding of IlvA’s allosteric regulation and offer opportunities for engineering feedback-resistant IlvA variants for biotechnological use.

## INTRODUCTION

Threonine dehydratase (IlvA; also known as threonine deaminase; EC 4.3.1.19) is the first step in the isoleucine arm of the branched-chain amino acid (BCAA) biosynthetic pathway (Fig. 1) (*1, 2*). The pyridoxal 5′-phosphate (PLP)-dependent reaction converts threonine to 2-iminobutyrate that is then converted by RidA (*3, 4*) to 2-ketobutyrate to continue isoleucine biosynthesis (*1, 2*). The BCAA pathway is regulated by IlvA and was one of the earliest examples of the first enzyme in a biosynthetic pathway being feedback regulated by the end-product, in this case isoleucine (Fig. 1) (*5, 6*). The kinetics of purified IlvA led Changeux to suggest that threonine and isoleucine bind to different sites on the enzyme, and that the binding of isoleucine decreases the binding of threonine (*7*). The ideas developed to explain IlvA kinetics led to the development of the two-state model that has become the blueprint for understanding the allosteric regulation of enzyme activity throughout biology (*8*).

**Figure 1.**
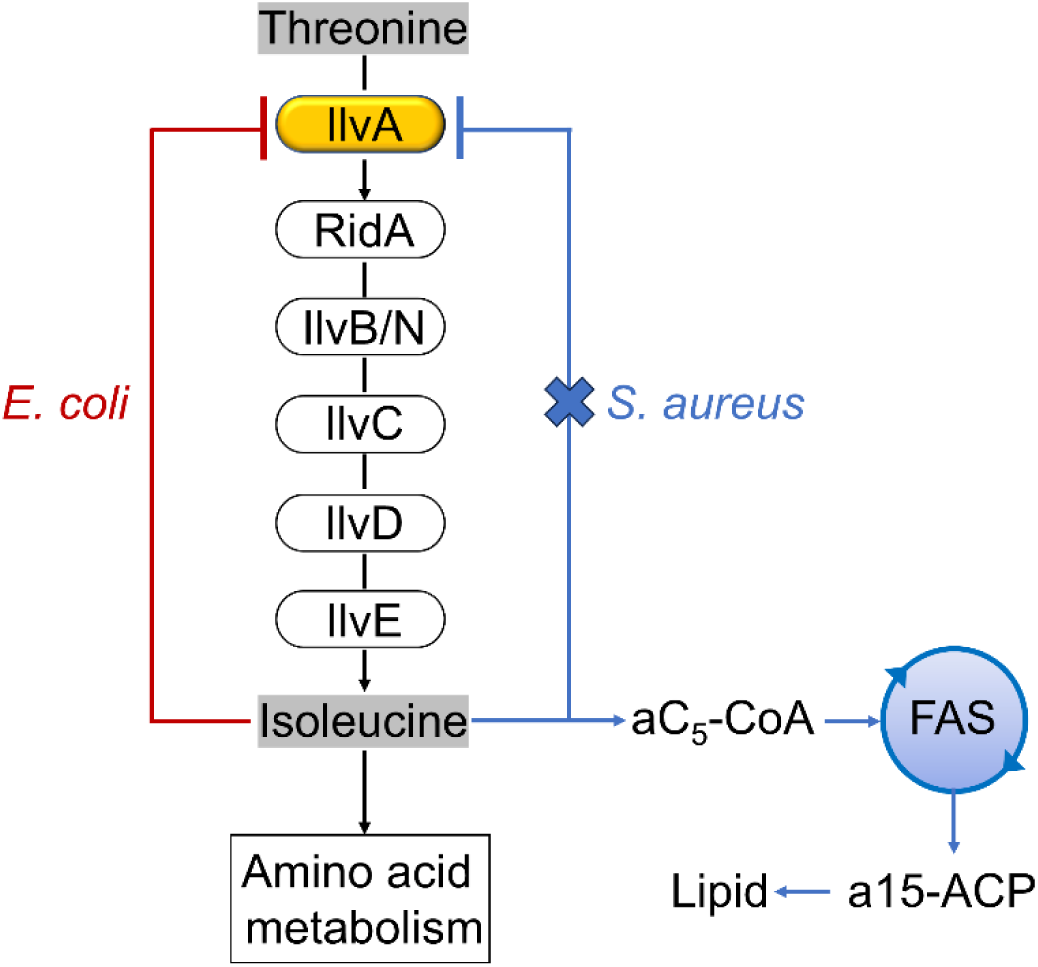
Role of threonine deaminase (IlvA) in the initiation of isoleucine biosynthesis. Threonine dehydratase/deaminase (IlvA) initiates the isoleucine arm of the branched-chain amino acid biosynthesis. RidA accelerates the spontaneous conversion of the IlvA product, 2-iminobutyrate, to 2-ketobuytrate, which is condensed with pyruvate by acetolactate synthase (IlvB/N) followed by ketoacid isomeroreductase (IlvC), dihydroxyacid dehydrase (IlvD) and branched-chain amino acid transaminase (IlvE) to result in isoleucine. Isoleucine is a potent feedback regulator of EcIlvA that maintains the intracellular isoleucine homeostasis by tuning the isoleucine biosynthetic rate to its utilization for protein synthesis. SaIlvA is not feedback regulated by isoleucine but is important for both amino acid metabolism and lipid metabolism in *S. aureus*.

BCAAs are essential for protein synthesis, but in *Staphylococcus aureus* they are also critical for fatty acid synthesis by generating necessary precursors (Fig. 1) (*9*). More specifically, isoleucine has a key role in *S. aureus* in controlling fatty acid composition and membrane stability/fluidity (*9–12*). The early studies on IlvA regulation were largely performed on the *Escherichia coli* enzyme (EcIlvA, UniProtKB P04968), but whether *S. aureus* IlvA (SaIlvA, UniProtKB Q2FF63) is regulated by isoleucine like EcIlvA is unknown. A comparison of the enzymes from the two organisms is important to further understand the regulatory panorama of this key enzyme, and this is the focus of the current study. Although much work has been performed on EcIlvA, including structure-function analyses, a number of key questions remain to be answered, and we also address these in the current study. A more complete understanding of the EcIlvA regulatory mechanisms is necessary for the comparison with SaIlvA, and how SaIlvA may also regulate fatty acid synthesis.

The previously determined 2.8 Å crystal structure of EcIlvA (PDB ID: 1TDJ) revealed that it is a tetramer, consisting of a dimer of carboxy-terminal regulatory domains projecting outward from a core formed by the tetramerization of the PLP-containing amino-terminal catalytic domains (*13, 14*). This EcIlvA structure appears to be the active (relaxed or R state) allosteric conformation of the assembly with the PLP cofactor covalently attached to Lys62 in the catalytic domain and isoleucine absent from the regulatory domain. Each EcIlvA carboxyl-terminal regulatory domain comprises two ACT-like subdomains (*13, 14*), a structural motif that is present in many regulatory enzymes involved in transcriptional control, amino acid and nucleotide metabolism (*15–17*). The prototypical ACT domain comprises 70-80 amino acids that fold into four β sheets and two α helices arranged in a βαββαβ structure. The first crystal structure containing an ACT domain was *E. coli* D-3-phosphoglycerate dehydrogenase (PGDH), a tetrameric protein with one ACT domain per subunit (*18*). The ACT domains form a symmetrical dimer, and two serine regulatory ligands were observed at the dimer interface. However, it has now been established that ACT domains can be combined in different ways to transmit allosteric regulatory signals through the binding of ligands at ACT domain interfaces (*15–17*). The isoleucine regulatory site in the EcIlvA regulatory domain has not been structurally characterized but has been inferred from molecular docking simulation studies using the crystal structure of the active EcIlvA coupled with kinetic analyses of site-directed mutations. These studies identified residue changes that result in resistance of IlvA to isoleucine inhibition, and these were interpreted as defining the isoleucine binding site (*19–21*). Finally, research has established that isoleucine is a potent IlvA inhibitor, whereas valine is not (*22–25*), but there is some disagreement as to whether there are two or four isoleucine binding sites in the *E. coli* IlvA (EcIlvA) (*19, 23, 26*).

Defining the structural transitions that switch EcIlvA from an active (R state) to an inactive (T state) conformation are key to understanding the mechanism of this important regulatory protein. Specifically, the identification of the isoleucine binding site and determination of how isoleucine binding triggers a conformational change that alters the EcIlvA active site environment remain key unanswered questions. Here, we report the structure of the isolated EcIlvA regulatory domain with isoleucine bound at a single site at the interface between the two ACT subdomains. The conformation of the isoleucine-bound regulatory domain is distinct from that of the isoleucine-free regulatory domain observed in the intact EcIlvA structure. We identify an allosteric pathway that transmits this information to the active site of an adjacent protomer in the tetrameric assembly. Having established how EcIlvA is regulated by isoleucine, we then investigated the regulation of SaIlvA and showed that it is refractory to isoleucine regulation. We discovered the structural basis for this observation by comparing the EcIlvA and SaIlvA regulatory domains, which revealed key differences in the ACT domain organization that leads to the disruption of the isoleucine binding site in SaIlvA. These conclusions are further supported by complementation and biochemical studies using an IlvA knockout strain of *S. aureus*.

## MATERIALS AND METHODS

### Materials

All chemicals and reagents were reagent grade. Pyridoxal phosphate (PLP) monohydrate was purchased from TCI (Product Number C0377). Crystallization screening kits and Sypro Orange dye were purchased from NeXtal Biotechnologies and Invitrogen, respectively. *E. coli* and *S. aureus* strains were routinely grown in rich broth (10 g/l tryptone, 5 g/l yeast extract, 5 g/l NaCl).

### Cloning, Expression and Purification of IlvA

The EcIlvA gene (b3772) and the regulatory domain (amino acids 335-514) of EcIlvA were amplified by PCR using primers designed for Gibson Assembly cloning into NdeI and EcoRI digested pET28a to obtain pPJ669 and pPJ668, respectively. Site directed mutagenesis was performed with pPJ669 and pPJ668 using quick change lightning mutagenesis (Stratagene) to obtain F352A mutant in the full length and regulatory domain of IlvA, respectively. SaIlvA (SaUSA300_2014) and its regulatory domain were amplified by PCR using primers designed for Gibson Assembly into pET28a. The SaIlvA was cloned into NcoI and XhoI digested vector while the regulatory domain was cloned into NdeI and EcoRI digested vector to obtain pPJ665 and pPJ666 respectively. The SaIlvA knockout (PDJ78) was generated in strain AH1263 by allelic replacement. For this approximately 1000 bp region the upstream and downstream of SAUSA300_2014 were PCR amplified using primers SaIlvA UP-EcoRI-F, SaIlvA UP-R, SaIlvA DN-F, SaIlvA DN-KpnI-R and gel purified. The PCR products were assembled into pJB38 by Gibson Assembly (New England Biolabs) yielding pPJ676. Plasmid pPJ676 was transformed into AH1263 by electroporation and the knockout was generated as previously described (*27*).

The knockout strain PDJ78 was confirmed by PCR. For testing complementation of ΔIlvA, the SaIlvA and EcIlvA genes were cloned into pCN38 vector (*28*). Briefly, for cloning EcIlvA, two fragments were generated by PCR, a 189 bp fragment containing the SarAP1 promoter, and 1633 bp fragment containing EcIlvA. Similarly, for SaIlvA, a 191 bp fragment containing the SarAP1 promoter and a 1347 bp fragment containing SaIlvA were generated. The two fragments for each EcIlvA and SaIlvA were assembled with SphI and BamHI digested pCN38 by Gibson Assembly. The plasmids were sequenced and transformed into ΔIlvA to test complementation. The pCN38 EcIlvA F352A mutant plasmid was generated by site-directed mutagenesis as described above. The list of primers and plasmids generated is given in Table S1. The pET28a plasmids containing the EcIlvA, EcIlvA(F352A), EcIlvA_R, EcIlvA_R(F352A), SaIlvA and SaIlvA_R were transformed into BL21(DE3) cells and grown to an OD600 of 0.7 at 37 °C with shaking at 200 rpm. The culture was induced for expression with 1mM IPTG and grown overnight at 16 °C with shaking at 200 rpm. For the full-length proteins EcIlvA, EcIlvA(F352A), and SaIlvA, cells were harvested and resuspended in 20 mM potassium phosphate buffer, pH 7.5, 500 mM NaCl, 0.1 mM TCEP, 1 mM PLP, 10 mM imidazole and were broken by two passages through a cell disruptor. The cell lysate was centrifuged at 20,000 g for 45 minutes and IlvA was purified from the supernatant using a Ni-NTA column by washing with 20 column volumes of each 20 mM potassium phosphate buffer, pH 7.5, 500 mM NaCl, 0.1 mM TCEP, 0.1 mM PLP containing 10 mM imidazole or 20 mM imidazole or 40 mM Imidazole. IlvA was eluted with 20 mM potassium phosphate buffer (pH 7.5), 500 mM NaCl, 0.1 mM TCEP, 0.1 mM PLP and 250 mM imidazole. The regulatory domain proteins IlvA_R and IlvA_R (F352A) pellets were resuspended in 20 mM Tris, pH 8.0, 500 mM NaCl, 10 mM imidazole. The protein was purified as described above except the buffer contained 20 mM Tris, pH 8.0, 500 mM NaCl and imidazole. SaIlvA_R pellets were resuspended in 20 mM Bis-Tris, pH 6.5, 500 mM NaCl, 10 mM imidazole, and 0.5% CHAPS. The protein was purified as described above except the buffer contained 20 mM Bis-Tris, pH 6.5, 500 mM NaCl, 250 mM imidazole and 0.5% CHAPS. All proteins except EcIlvA_R were further purified by gel filtration chromatography using a HiLoad 16/600 Superdex 200 column (Cytiva Life Sciences). EcIlvA_R was dialyzed against 20 mM Tris, pH 7.5. 200 mM NaCl. EcIlvA, and EcIlvA(F352A) were eluted with 20 mM potassium phosphate buffer, pH 7.5, 200 mM NaCl, 0.1 mM TCEP. EcIlvA_R(F352A) was eluted with 20 mM Tris, pH 7.5, 200 mM NaCl. SaIlvA was eluted with 20 mM potassium phosphate buffer, pH 7.5, 500 mM NaCl, 0.1 mM TCEP. SaIlvA_R was eluted with 20 mM Bis-Tris, pH 6.5, 200 mM NaCl. The molecular weight was calculated by analyzing the peak size on a HiLoad 13/300 Super 200 analytical column (Cytiva Life Sciences). A standard curve was generated using thyroglobulin (669 kDa), ferritin (440 kDa), aldolase (158 kDa), conalbumin (75 kDa), ovalbumin (44 kDa), ribonuclease A (13.7 kDa), and aprotinin (6.5 kDa).

### IlvA activity assay

The activity assay was performed by a spectrophotometric method using purified recombinant EcIlvA, EcIlvA(F352A) and SaIlvA proteins as described previously (*19, 20, 29*). Briefly, the reaction mixture contained 100 mM potassium phosphate buffer, pH 7.5, 20 μM PLP, 50 nM enzyme, and varied concentration of threonine (0 to 100 mM) for EcIlvA and EcIlvA(F352A). Assays were carried out in the presence and absence of 0.05 mM and 0.5 mM isoleucine for EcIlvA and 5 mM isoleucine for EcIlvA(F352). The effect of isoleucine on the activities of EcIlvA and EcIlvA(F352A) were assayed at the presence of 7 mM threonine and varied concentration of isoleucine (0 to 0.5 mM). The reaction mixture was the same except 200 nM of enzyme and varied concentration of threonine (0 to 200 mM) for SaIlvA. Assays for SaIlvA were carried out in the presence and absence of 0.5 mM and 5 mM isoleucine. The effect of isoleucine on the activity of SaIlvA was assayed at the presence of 30 mM threonine and varied concentration of isoleucine (0 to 5 mM). The reaction was carried at 30 °C and the formation of 2-ketobutyrate was measured continuously at 230 nm on a 96-well clear flat bottom UV-transparent microplate (Corning, Product No. 3635) using a BioTek Synergy H1 Microplate Reader (BioTek Instruments, Inc.). All measurements were repeated three times. The enzyme kinetics were calculated by fitting allosteric sigmoidal and Michaelis-Meneten equations for EcIlvA and SaIlvA, respectively, within GraphPad Prism 10.0.2 (GraphPad Software, Boston, Massachusetts USA, www.graphpad.com).

### X-ray crystallography

EcIlvA, EcIlvA_R•Ile, EcIlvA_R(F352A)•Ile, and SaIlvA_R crystals were grown by the sitting and hanging drop vapor diffusion methods at 20 °C using a Mosquito crystallization robot (TTP Labtech). A final 5 mM isoleucine was added to the EcIlvA protein solution (10.2 mg/ml in 20 mM potassium phosphate, pH 7.5, 0.2 M sodium chloride, and 0.1 mM TCEP) and incubated for 30 minutes at room temperature before crystallization. The sitting drops were produced by mixing 0.2 μl of the protein solution with 0.2 μl of reservoir solution (0.1 M HEPES, pH 7.0, 0.1 M magnesium chloride, and 15% PEG 4000). Crystals were transferred into a cryo-protection solution (0.1 M HEPES, pH 7.0, 0.1 M magnesium chloride, 15% PEG 4000, 1 mM isoleucine, and 30 % glycerol) and flash frozen in liquid nitrogen. Diffraction data were collected at the SERCAT beamline 22-ID at the Advanced Photon Source and processed using xia2 (*30*). The EcIlvA structure was solved by molecular replacement method with EcIlvA molecule (PDB ID: 1TDJ) as a search model using Phaser Crystallographic Software (*31*). To obtain the EcIlvA_R•Ile crystals, the sitting drops were produced by mixing 0.2 μl of the purified EcIlvA_R protein solution (10 mg/ml in 20 mM Tris-HCl, pH 7.5, 0.2 M sodium chloride, and 0.1 mM EDTA) with 0.2 μl of reservoir solution (0.2 M ammonium acetate and 20% PEG 3350). For cryoprotection, crystals were transferred into a reservoir solution supplemented with 30% glycerol, and subsequently cryocooled in liquid nitrogen. X-ray diffraction data were collected at the 17-ID-1 AMX beamline (*32*) at the National Synchrotron Light Source II (NSLS II), Brookhaven National Laboratory. Diffraction data were processed using autoPROC toolbox (*33*) coupled with XDS (*34*), POINTLESS (*35*), and AIMLESS (*35*). The EcIlvA_R structure was solved by molecular replacement method with the regulatory domain (residues 335-514) of the EcIlvA molecule (PDB ID: 1TDJ) as a search model using Phaser Crystallographic Software. To obtain the EcIlvA_R(F352A)•Ile crystals, a final 5 mM isoleucine was added to the EcIlvA_R(F352A) protein solution (9.4 mg/ml in 20 mM Tris-HCl, pH 7.5, and 0.2 M sodium chloride). Crystals were grown using a reservoir consisting of 0.04 M sodium acetate, pH 4.0, 0.08 M ammonium acetate, and 6% PEG 4000. Crystals were transferred into a cryo-protection solution (0.07 M sodium acetate, pH 4.0, 0.14 M ammonium acetate, 10.5% PEG 4000, 2 mM isoleucine, and 30 % glycerol) and flash frozen in liquid nitrogen. Diffraction data were collected at beamline 5.0.2 of the Advanced Light Source. Diffraction data were processed using XDS and AIMLESS. The EcIlvA_R(F352A)•Ile structure was solved by molecular replacement method with EcIlvA_R•Ile structure (PDB ID: 9D2R) as a search model using Phaser Crystallographic Software. To obtain the SaIlvA_R crystals, the hanging drops were produced by mixing 1 μl of the purified SaIlvA_R protein solution (10 mg/ml in 20 mM Bis-Tris, pH 6.5, and 0.2 M sodium chloride) with 1 μl of reservoir solution (0.1 M CAPSO, pH 9.0, 1.35 M ammonium sulfate and 0.2 M sodium chloride). For cryoprotection, crystals were transferred into a mixture of mineral oil and Paraton-N, and flash frozen in liquid nitrogen. Diffraction data were collected at the SERCAT beamline 22-ID at the Advanced Photon Source and processed using HKL2000 (*36*). The SaIlvA_R structure was solved by molecular replacement method with the regulatory domain (residues 336-422) of the SaIlvA AlphaFold model structure (AF-Q2FF63) as a search model using Phaser Crystallographic Software. All structures were refined and optimized using PHENIX (*37*) and COOT (*38*), respectively. Data collection and refinement statistics are summarized in Table S2. Figures were rendered with PyMOL (version 2.5.5, Schrödinger, LLC).

### SEC-SAXS analysis

SEC-SAXS was performed at BioCAT (beamline 18ID at the Advanced Photon Source, Chicago) with in-line size exclusion chromatography (SEC) to separate sample from aggregates and other contaminants thus ensuring optimal sample quality and multiangle light scattering (MALS), dynamic light scattering (DLS) and refractive index measurement (RI) for additional biophysical characterization (SEC-MALS-SAXS). The samples were loaded on a Superdex 200 Increase 10/300 GL column (Cytiva) run by a 1260 Infinity II HPLC (Agilent Technologies) at 0.6 ml/min. The flow passed through (in order) the Agilent UV detector, a MALS detector and a DLS detector (DAWN Helios II, Wyatt Technologies), and an RI detector (Optilab T-rEX, Wyatt). The SAXS sheath flow cell (*39*) consists of a 1.0 mm internal diameter quartz capillary with ∼20 mm walls (effective path length 0.542 mm). A coflowing buffer sheath is used to separate samples from the capillary walls to minimize radiation damage. Scattering intensity was recorded using an Eiger2 XE 9M (Dectris) detector which was placed 3.6 m from the sample giving us access to a q-range of 0.003 Å-1 to 0.42 Å-1. Exposures (0.5 s) were acquired every 1 s during elution. Buffer blanks were created by averaging regions flanking the elution peak and subtracted from exposures selected from the elution peak to create the I(q) vs q curves used for subsequent analyses. SAXS data were processed, and the overall parameters computed following standard procedures of the software package BioXTAS RAW 2.1.4 (*40*). The Guinier plots, the P(r) function and the SAXS parameters in Table S3 were calculated using described methods (*41–45*). The Evolving factor analysis (EFA) function (*46*) in RAW for EcIlvA_R was used to deconvolve overlapping chromatography data. The intensity at zero scattering angle I(0) and radius of gyration Rg were obtained by performing a linear fit to the Guinier plot. The particle dimension Dmax was determined from the pair-distance distribution P(r) with the program GNOM (*41*). Reconstructions (n = 20) were aligned, averaged, and refined in slow mode using DENSS (*47*) to calculate the electron density for the final three-dimensional volume. For EcIlvA and EcIlvA_R, the ‘Align output to PDB/MRC’ option was used with EcIlvA tetramer model (PDBID: 9D2Q) and EcIlvA_R dimer model (PDBID: 9D2R), respectively, during the DENSS run. The EcIlvA tetramer model, the EcIlvA_R dimer model and SaIlvA_R octamer model were fit into their SAXS electron density using UCSF ChimeraX, version 1.8 (*48*). Theoretical scattering profiles and qualities of fit (χ2) were obtained from the FoXS webserver (*49*) using the atomic coordinates of EcIlvA tetramer, EcIlvA_R dimer, SaIlvA octamer and their experimental SAXS profiles. The theoretical dimensionless Kratky plots were computed in RAW using the theoretical scattering profiles obtained from FoXS.

### Thermal stability measurements

Purified proteins EcIlvA, EcIlvA(F352A), EcIlvA_R, EcIlvA_R(F352A) and SaIlvA were subjected to thermal denaturation in the presence of Sypro Orange dye as described in (*50*). Briefly, 10 μM of each of the proteins were added to 30 μl of 20 mM HEPES, pH 7.5, 150 mM NaCl and 10X Sypro Orange dye in a 96 well plate. The stability of the proteins was analyzed in the presence and absence of 5 mM isoleucine. The plate was centrifuged at 1500 rpm for 2 min before being placed in an Applied Biosciences 7500 RealTime PCR instrument. A thermal scan from 25 °C to 95 °C was performed using an increment rate of 1 °C/min. All experiments were performed in triplicate. The data were fitted using Boltzmann sigmoidal equation within GraphPad Prism 10.0.2 (GraphPad Software, Boston, Massachusetts USA, www.graphpad.com).

### Molecular docking

Molecular docking of threonine substrate with the EcIlvA protein was performed using Molecular Operating Environment (MOE) software (2018.01; Chemical Computing Group). For docking calculation, the threonine ligand was drawn using ChemDraw Professional and converted to a 3D model (version 17.1, PerkinElmer Informatics). An IlvA-PLP structure for docking calculation was generated from the IlvA structure (PDB ID: 9D2Q). All molecules except the protein and PLP were removed. The ligand and protein structures were imported into MOE and prepared using the “QuickPrep” function. After performing docking using the “Dock” function of MOE, a docking model with the possibility of reacting to the cofactor PLP was selected.

### EcIlvA and SaIlvA assay using radioactive threonine

5 mM 14[C] threonine-specific activity (989 µCi/mmol) was incubated with 5 μM EcIlvA or SaIlvA in the presence of 40 μM PLP and 50 mM HEPES buffer or phosphate buffer (pH 7.5) for 30 minutes. 10 μl of the assay was spotted on a Silica gel 60 thin layer chromatography plate. The TLC was resolved using 1-Butanol: acetic acid: water (80:20:4 v/v/v). The TLC was air-dried and exposed to a phosphorimager screen overnight and scanned using Typhoon FLA9500 (GE Healthcare)

### Growth curve and fatty acid analysis

For growth curve and fatty acid analysis ΔIlvA strain containing empty vector (pCN38) or pSaIlvA or pEcIlvA grown in Complete Defined Medium (CDM) overnight. This was diluted to OD600 of 0.1 and grown in CDM or CDM minus isoleucine for 5-6 hours. For fatty acid analysis the cells were harvested after 6 hours and lipids were extracted by Bligh Dyer method (*51*). Methyl esters of the fatty acids were generated with the lipid fraction and analyzed by gas chromatography (*52*).

### EcIlvA and SaIlvA spectra

The spectral scan of EcIlvA and SaIlvA was performed using nanodrop in the presence of Buffer (20 mM K PO4 Buffer (pH 7.5), 200 mM NaCl, 0.1 mM TCEP) plus 0.1 mM PLP, or 30 mM threonine as indicated in the figure. The data was smoothed using Graphpad prism software.

## RESULTS

### Crystal structure of *E. coli* IlvA

To facilitate the analysis of the EcIlvA allosteric regulatory mechanism, we first attempted to obtain a higher resolution crystal structure of the enzyme than the current 2.8 Å structure (PDB ID: 1TDJ) (*13*). His-tagged EcIlvA (Fig. S1*A*) was purified by affinity chromatography followed by gel filtration chromatography (Fig. S1*B*), and we obtained crystals in space group I222. The hexa-His tag was not removed, and His tagged protein was used for all the assays. The structure was determined at 1.87 Å resolution, and the asymmetric unit contains one EcIlvA monomer that is composed of two globular domains (Fig. 2*A* and Table S2). The amino terminal catalytic domain comprises residues 1-320 (Fig. S2*A*) and contains the enzyme active site with PLP covalently bound to Lys62 (Figs. 2*A* and S2*B*), and the carboxy terminal regulatory domain comprises residues 335-514 (Figs. 2*A* and S2*B*). Residues 482-496 in the regulatory domain are disordered and their conformation could not be determined (Fig. S2*B*). The independently-folded catalytic and regulatory domains are linked by helix H14 that comprises residues 321-334 (Fig. S2*A*), which is referred to as the ‘neck’ (*13*) (Figs. 2*A* and S2*B*). Overall, the higher resolution EcIlvA structure superimposed well with the published structure (PDB ID: 1TDJ) with the root mean square deviation (RMSD) of 1.4 Å over 493 Cα atoms. Comparing this higher resolution EcIlvA structure with the published structure, we found that four regions had different conformations (Fig. 2*A*). Inspection of the published structure with the electron density maps reveals that minor corrections are needed in the misfitted regions (regions 1 to 3) of the published structure (Fig. S3). Region 1 is close to the catalytic site and region 3 (loop L22) is involved in the conformational coupling of the regulatory and catalytic domains (see below). The corrected models of these regions exhibit similar conformations to the higher resolution structure (Fig. S3). The conformational difference in region 4 is related to the binding of glycerol. Consequently, the higher resolution glycerol-bound EcIlvA structure provides higher quality structural information.

**Figure 2.**
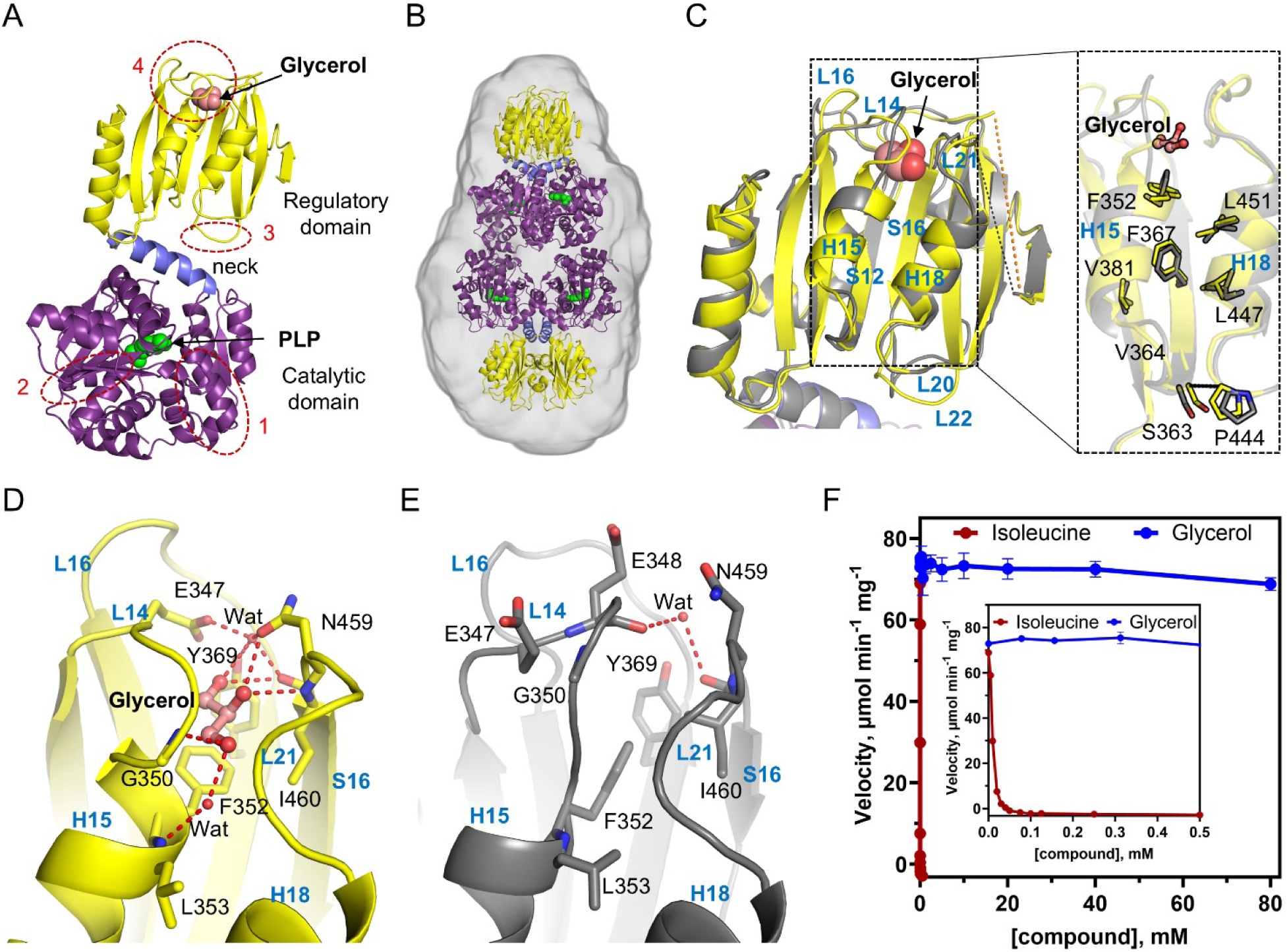
Crystal structures of *E. coli* IlvA. *A*, structure of the EcIlvA monomer at 1.87 Å resolution (PDB ID: 9D2Q). It consists of an amino terminal catalytic domain (purple) connected to a carboxy terminal regulatory domain (yellow) by helix H14 (the neck, blue). This new higher resolution EcIlvA structure clarifies the conformation of 3 regions in the previously reported structure (PDB ID: 1TDJ) (dashed red circles, 1 to 3). The PLP cofactor (green balls) is covalently attached to Lys62 in the catalytic domain and there is a glycerol (salmon balls) bound in the regulatory domain. *B*, the aligned electron density map occupied by EcIlvA in solution was determined by small angle X-ray scattering (SAXS) and matches the crystal structure of the tetramer. *C*, the EcIlvA regulatory domain in the new structure (PDB ID; 9D2Q) (yellow) overlaid with the regulatory domain in (PDB ID: 1TDJ) (gray) illustrating how the conformation of loops L14, L16 and L21 are altered by glycerol binding. *Inset*, a close-up view of the area where the conformational change occurs upon binding of isoleucine. *D*, the regulatory domain conformational change that occurs within the pocket in the presence of bound glycerol. Glycerol is depicted as salmon balls-sticks, hydrogen bond interactions are shown with dotted red lines, and water molecules are red spheres. *E*, the glycerol binding pocket in the ligand-free EcIlvA regulatory domain (PDB ID: 1TDJ). A water molecule is shown with red sphere. *F*, the dose response curve shows the effect of glycerol on the activity of EcIlvA. The initial velocity was measured in the presence of 7 mM threonine. Isoleucine was used as the inhibitor control compound.

The EcIlvA tetramer is organized as a dimer of dimers, and the protomers are identical in our structure because the tetramer is generated through crystallographic symmetry operations (Fig. S2*C*). Gel filtration chromatography (Fig. S1*C*) and prior work (*13*) are consistent with the observed tetrameric organization of the EcIlvA protomers. The amino terminal catalytic domains create the core tetramer with PLP bound to Lys62 in all four protomers (Fig. S2*C*), and the carboxy terminal regulatory domains form dimers that protrude from either side of the core. To validate the crystal structure organization, we used size-exclusion chromatography coupled with small angle X-ray scattering (SEC-SAXS) analysis to compare the symmetry generated EcIlvA tetramer with the volume occupied by the EcIlvA tetramer in solution (Fig. S4). SAXS experiments showed that the sample was globular and monodisperse tetramer with a molecular weight of 239.6 kDa (Figs. S4*A*, S4*D*, and Table S3) (*53*). A comparison of the EcIlvA SAXS experimental data set with the predicted profile of the EcIlvA tetramer based on the crystal structure (Fig. S2*C*) shows that the SAXS data are a very close fit to the theoretical behavior of the EcIlvA tetramer (χ^2^ = 1.34) (Fig. S4*G*). The EcIlvA tetramer fits very well into the aligned *ab initio* electron density maps generated from the solution scattering data, with the correlation coefficient of 0.92. This shows a close agreement between the EcIlvA crystal and solution structures (Fig. 2*B*). SAXS data indicates that the crystallographic organization is present in solution.

The regulatory domain contains a bound glycerol in our structure that was not present in PDBID: 1TDJ (Figs. 2*A* and S2*B*). This difference in the two structures likely arises from the use of glycerol as a cryoprotectant in our structure, whereas the prior analysis did not use frozen crystals (*13, 14*). The bound glycerol appears to elicit a conformational change in the regulatory domain. The glycerol and two structured waters are bound by helix H15, loops L14 and L21 that cluster at the tip of the regulatory domain furthest from the catalytic domain (Fig. 2*C*). The 2Fo-Fc electron density map clearly shows the bound glycerol and the two structured waters (Fig. S5*A*). The hydroxyl groups of glycerol form hydrogen-bonding interactions with the main chain carbonyl oxygen atom of Ile460 and amide nitrogen atoms of Gly350 and Ile460, with the amide nitrogen atom of Leu353 and side chains of Glu347 and Tyr369 via water molecules (Fig. 2*D*). The 3 carbon atoms of glycerol engage in a hydrophobic pocket composed of residues Phe352 and Ile460. This glycerol binding pocket is significant because it corresponds to the isoleucine binding site as discussed below. In PDB ID: 1TDJ, residues Glu347 and Asn459 are pointing toward the solvent, and a water molecule interacts with the main chain carbonyl oxygen atoms of residue Glu348 on loop L14 and Ile460 on loop L21 (Fig. 2*E*). Comparing the regulatory domains of the glycerol-free and glycerol-bound structures, there are structural differences in Phe352 and Loop L14, but the conformation of Phe367 and its adjacent residues remain unchanged (Fig. 2*C*). The dose-response curve indicates that isoleucine effectively inhibits EcIlvA activity, whereas glycerol, despite binding to the isoleucine binding site of regulatory domain, shows negligible inhibition activity, maintaining an activity level of 93.4% even at a high concentration of 80 mM (Fig. 2*F*). Therefore, both glycerol-free and glycerol-bound EcIlvA were used as the isoleucine-free EcIlvA structure for further discussion.

### Crystal structure of the *E. coli* IlvA regulatory domain (EcIlvA_R)

We failed to obtain full-length EcIlvA crystals containing bound isoleucine. Therefore, we engineered an expression construct that encompassed the proposed EcIlvA regulatory domain starting at residue Glu335 (EcIlvA_R) based on the EcIlvA crystal structure (Figs. S1*A* and S2*A*). The isolated EcIlvA_R was soluble and successfully purified (Fig. S1*B*), and was a 36 kDa dimer based on gel filtration chromatography (Fig. S1*C*). Crystals were obtained in space group P3_2_21, and the structure was determined at 1.60 Å resolution (Table S2). There is one EcIlvA_R in the asymmetric unit (Fig. 3*A*) and the dimeric assembly is generated by crystallographic symmetry (Fig. 3*B*). Each EcIlvA_R comprises six helices (H15-H20) and eight strands (S11-S18) (Fig. 3*A*) folded into two ACT-like subdomains connected by loop L19: ACT1 (helices H15-H17 and strands S11-S14) and ACT2 (helices H18-H20 and strands S15-S18). ACT1 and ACT2 are very similar with the RMSD of 2.3 Å over 69 Cα atoms. The prototypical ACT domain has a βαββαβ topology but ACT1 and ACT2 each have an extra helix (H17 and H20, respectively) leading to a βαββαβα organization (Fig. 3*A*). H17 and H20 form the dimerization interface (Fig. 3*B*). The disordered region (482–496) in the regulatory domain of the full-length EcIlvA structure was well-defined in the EcIlvA_R structure and comprises loop L23 and helix H19 (Figs. 3D and S2*A*). To validate crystal structure organization, we used SEC-SAXS analysis to compare the dimeric EcIlvA_R crystal structure with the volume occupied by the EcIlvA_R dimer in solution (Figs. S4). SAXS experiments showed that EcIlvA_R behaves as a dimer with molecular weight of 42.4 kDa (Figs. S4*B*, S4*E* and Table S3). A comparison of the EcIlvA_R experimental data with the predicted profile of the EcIlvA_R dimer based on the crystal structure (Fig. 3*B*) shows that the SAXS data are a very close fit to the theoretical behavior of the EcIlvA_R dimer (χ^2^ =3.05) (Fig. S4*H*). The EcIlvA_R crystallographic dimer was fitted into the aligned *ab initio* electron density map generated from the solution scattering data, with the correlation coefficient of 0.82. This shows the close alignment of the EcIlvA_R crystal and solution structures (Fig. 3*C*).

**Figure 3.**
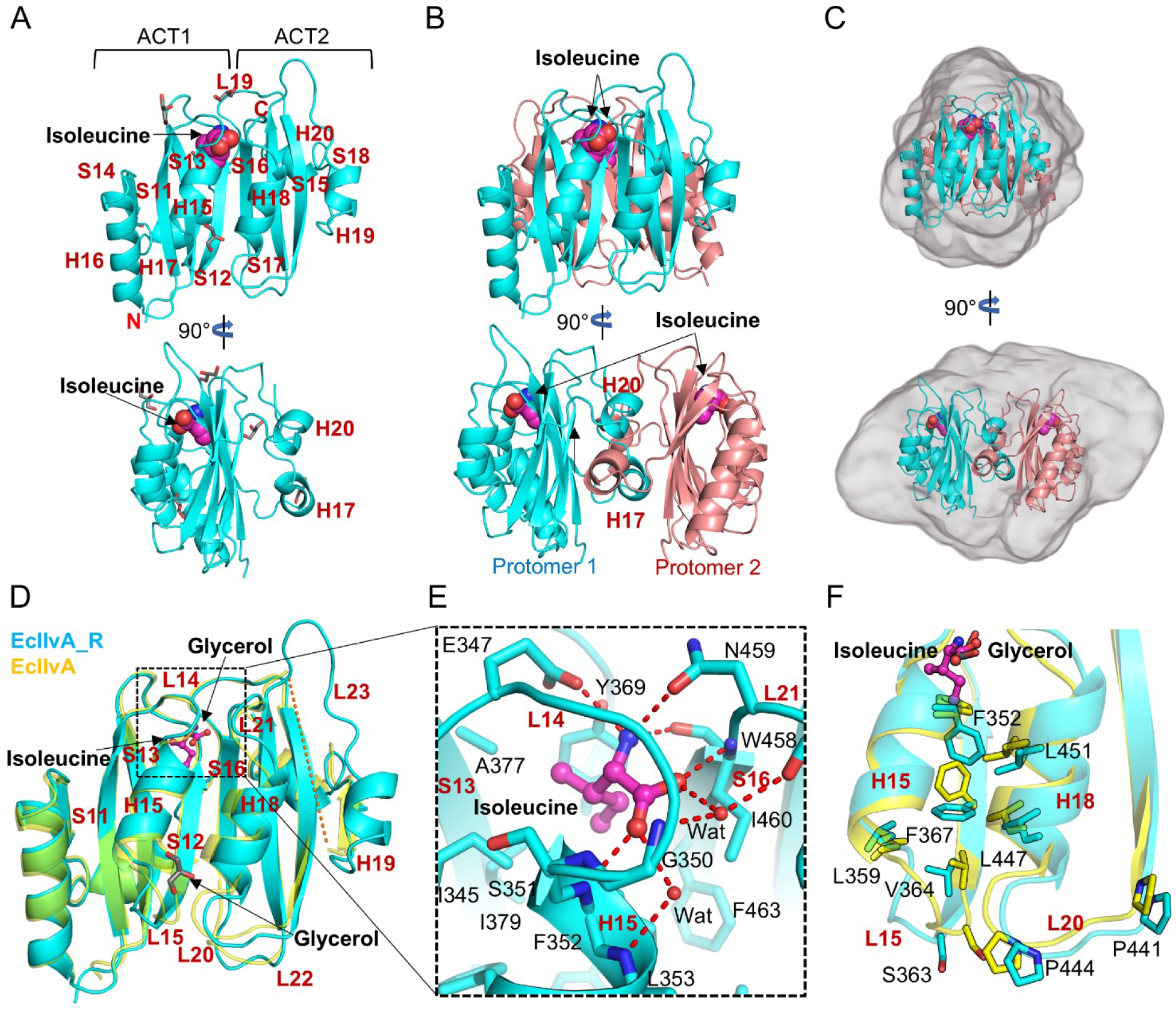
Crystal structure of the EcIlvA_R•Ile complex. *A*, overview of the EcIlvA_R•Ile complex protomer with one isoleucine molecule (magenta balls) bound (PDB ID: 9D2R). The rotated view (below) illustrates the positions of the two helices (H17 and H20) that are added to the core structure consisting of two ACT domains inked by loop L19. Five glycerol molecules are depicted as gray sticks. *B*, the structure of dimeric EcIlvA_R•Ile complex with the protomers shown in cyan and salmon. The rotated view (below) shows how helices H17 and H20 mediate the formation of the dimer. *C*, the EcIlvA_R•Ile dimer fits well into the aligned electron density experimentally determined by SAXS in solution. *D*, superimposition of EcIlvA_R•Ile (cyan) with the glycerol (salmon balls-sticks) bound regulatory domain of EcIlvA (yellow) (PDB ID: 9D2Q). Isoleucine (magenta balls-sticks) is bound between loops L14 and L21 triggering structural changes in the two adjacent helices H15 and H18 and loops L15, L20, and L22. A glycerol molecule, depicted as gray sticks, is bound around Loop L15 of EcIlvA_R•Ile. *E*, close-up views of the isoleucine binding site. Isoleucine is depicted as magenta balls-sticks, water molecules (Wat) are red balls and hydrogen bonds are dashed red lines. *F*, an overlay of the EcIlvA_R•Ile complex (cyan) with the EcIlvA regulatory domain (yellow; PDB ID: 9D2Q). The change in the sidechain rotamer of Phe352 must occur to accommodate isoleucine and leads to a cascade of structural changes along helices H15 and H18 that result in the movement of loops L15 and L20.

The EcIlvA_R structure revealed the presence of one isoleucine molecule (Figs. 3*A* and S5*B*) that binds between sheets S11-S13, loop L14 and helix H15 of the ACT1 subdomain, and loop L21 and sheet S16 of ACT2 subdomain (Fig. 3*D*). This is the same location where glycerol is bound to full-length EcIlvA (Fig. *3D*). The amino group of isoleucine forms hydrogen-bonding interactions with the main chain carbonyl oxygen atom of Ile460 and side chains of Glu347 and Asn459 that close over the isoleucine pocket (Fig. 3*E*). The isoleucine carboxyl group forms hydrogen-bonding interactions with the amide nitrogen atoms of Phe352 and Ile460. Residues Ser351 and Phe352 are at the amino terminus of helix H15, which also allows a favorable dipole interaction with the carboxyl group. In addition, the carboxyl group interacts with the carbonyl oxygen atom of Trp458 and the amide nitrogen atoms of Gly350 and Leu353 via water molecules (Fig. 3*E*). The side chain of the bound isoleucine engages a hydrophobic pocket composed of residues Ile345, Ser351, F352, Tyr369, Ala377, Ile379, Ile460, and Phe463 (Fig. 3*E*).

Notably, there is a conformational change in EcIlvA_R (Fig. 3*F*) that distinguishes it from the isoleucine-free regulatory domain of EcIlvA (Fig. 2*C*). The side chain of Phe352 rotates downward to accommodate the isoleucine terminal methyl group, which triggers a series of structural changes in the residues along helices H15 and H18 (F352-L451-F367-L447-V364) that result in the movement of loops L15 and L20 some 23 Å away from the isoleucine pocket (Fig. *3F*). Loop L20 places the Cα atom of Pro444 3 Å away from its position in the isoleucine-free structure. There are a total of five glycerol molecules in the EcIlvA_R structure, two of them bound in the dimer interface (Fig. 3*A*). Interestingly, a glycerol molecule is bound in a pocket around loop L15, which is formed by a conformational change due to isoleucine binding (Figs. 3*A* and 3*D*). In the isoleucine-free and isoleucine-bound structures, the distances between the Cα atoms of Ser363 and Pro444 are 5.0 Å and 8.6 Å, respectively. Loops L15 and L20 are further apart in the isoleucine-bound structure (Fig. 3*F*). These data show that isoleucine binding requires the movement of the Phe352 sidechain that, in turn, initiates a conformational change that ripples across the EcIlvA regulatory domain.

### Biochemical analysis of EcIlvA(F352A)

The structural analysis of EcIlvA_R indicates that Phe352 is the key residue in the isoleucine binding site that is required to initiate the conformational signal when the ligand binds. Therefore, we purified and characterized the EcIlvA(F352A) mutant (Figs. S1*A*-*C*). As reported (*13, 22–25*), kinetic analysis shows that isoleucine inhibits the binding of threonine to EcIlvA (Fig. 4*A*). In the enzymatic assays used here, the formation of the IlvA product 2-ketobutyrate was measured (*19, 20, 29*) and hence the aldimine formation between threonine and PLP was not used for as a readout for the enzymatic assay. Threonine binding is normally cooperative (n_H_ = 1.9) and at 0.5 mM, isoleucine extinguishes EcIlvA activity regardless of the threonine concentration (Figs. 4*A* and 4*C*). Threonine binding to EcIlvA(F352A) has approximately the same binding kinetics as to wild type EcIlvA, but the binding to EcIlvA(F352A) is not cooperative (n_H_ = 1.04) and the enzyme remained active in the presence of 5 mM isoleucine (Figs. 4*B* and 4*C*). The ability of isoleucine to inhibit EcIlvA and EcIlvA(F352A) was directly compared, and isoleucine is a potent inhibitor of EcIlvA, but not EcIlvA(F352A) (Figs. 2*F* and 4*D*). Thus, the kinetic analysis of EcIlvA(F352A) supports the proposed role of Phe352 in initiating the conformation change required for isoleucine to inhibit EcIlvA. It was essential to demonstrate that the inability of isoleucine to inhibit the EcIlvA(F352A) mutant was not simply due to the lack of binding to the enzyme. We therefore used a thermal stabilization assay to measure isoleucine binding to EcIlvA and EcIlvA(F352A) (Fig. *4E*). EcIlvA and EcIlvA(F352A) had nearly identical stabilities to thermal denaturation in the absence of isoleucine, and the stability of both proteins was increased ∼14 °C in the presence of isoleucine. These data confirm that both proteins bind isoleucine (Fig. *4E*).

**Figure 4.**
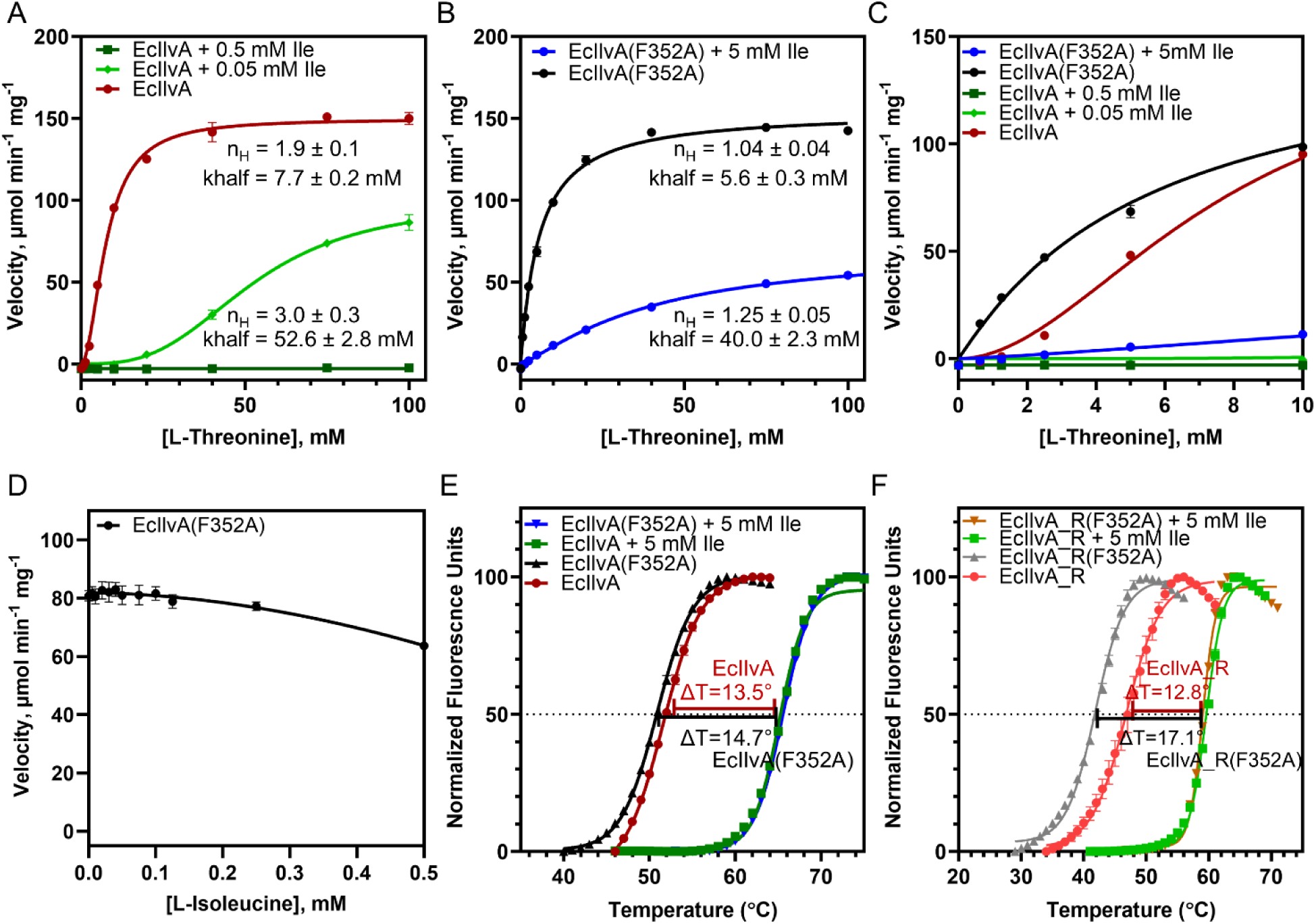
Isoleucine effect on the activity and stability of EcIlvA. A, effect of isoleucine on the steady-state kinetics of EcIlvA. *B*, effect of isoleucine on the steady-state kinetics of EcIlvA(F352A). The theoretical curve was analyzed by the allosteric sigmoidal equation using GraphPad Prism 9.4.1. for both EcIlvA and EcIlvA(F352A). The kcat values for EcIlvA and EcIlvA(F352A) were calculated to be 144.7 ± 1.5 sec^-1^ and 149.2 ± 2.2 sec^-1^, respectively. Khalf, kcat and Hill coefficient (n_H_) values are presented as mean values ± standard error (SE). *C*, close-up plots showing the steady-state kinetics of EcIlvA and EcIlvA(F352A) at the low concentration of threonine (0 – 10 mM). *D*, effect of isoleucine on the activity of EcIlvA(F352A). The initial velocity was measured in the presence of 7 mM threonine. *E*, the thermal stability of EcIlvA and EcIlvA(F352A) in the absence and presence of 5 mM isoleucine. *F*, the thermal stability of EcIlvA_R and EcIlvA_R(F352A) in the absence and presence of 5 mM isoleucine. Data represented as mean ± standard deviation (error bars) from triplicate sets.

### Crystal structure of the EcIlvA regulatory domain F352 mutant in complex with isoleucine (EcIlvA_R(F352A)•Ile)

The impact of the Phe352 mutation on the regulatory domain conformation was assessed by purifying EcIlvA_R(F352A) (Figs. S1*A*-*C*) and solving the crystal structure of the EcIlvA_R(F352A)•Ile complex (Table S2 and Fig. S5*C*). The EcIlvA_R(F352A) protein was less stable by 4.9 °C to thermal denaturation compared to EcIlvA_R, but the presence of isoleucine greatly stabilized both proteins illustrating that isoleucine binds to EcIlvA_R(F352A) (Fig. *4F*). These results using the isolated regulatory domain agree with the data for the full-length protein (Fig. *4E*). Crystals of the EcIlvA_R(F352A)•Ile complex were obtained in space group C2, and the structure was determined at 1.22 Å resolution (Table S2). The asymmetric unit contains a monomer, and the dimeric assembly was generated through crystallographic symmetry. Like the structure of EcIlvA_R•Ile, an isoleucine molecule is bound at the same location in EcIlvA_R(F352A)•Ile (Figs. 5*A* and S5*C*). However, due to the absence of the Phe352 sidechain that initiates the conformational change in the regulatory domain (Fig. 3*F*), isoleucine binding does not trigger a conformational change in the EcIlvA_R(F352A) mutant (Fig. 5*A*). The overlay of EcIlvA_R(F352A) with the regulatory domain of EcIlvA shows that the two structures have the same conformation. A close-up view of the residues between the helices H15 and H18 shows that the sidechains of Phe355, Leu359, Val364, Phe367, Leu447 and Leu451 in EcIlvA_R(F352A) all adopt the same conformation as found in the Isoleucine-free (glycerol bound) regulatory domain of EcIlvA (Fig. 5*B*). Loops L15 and L20 are superimposed, and the Cα distance between Ser363 and Pro444 is 4.4 Å, which is similar to the Isoleucine-free regulatory domain of EcIlvA. A comparison of the overall structure of EcIlvA_R(F352A)•Ile to EcIlvA_R•Ile highlights their differences (Fig. 5*C*), and a close-up view of helices H15 and H18 illustrates how the absence of the Phe352 sidechain prevents the conformational change caused by isoleucine binding to extend down the regulatory domain (Figs. 5*B* and 5*C*).

**Figure 5.**
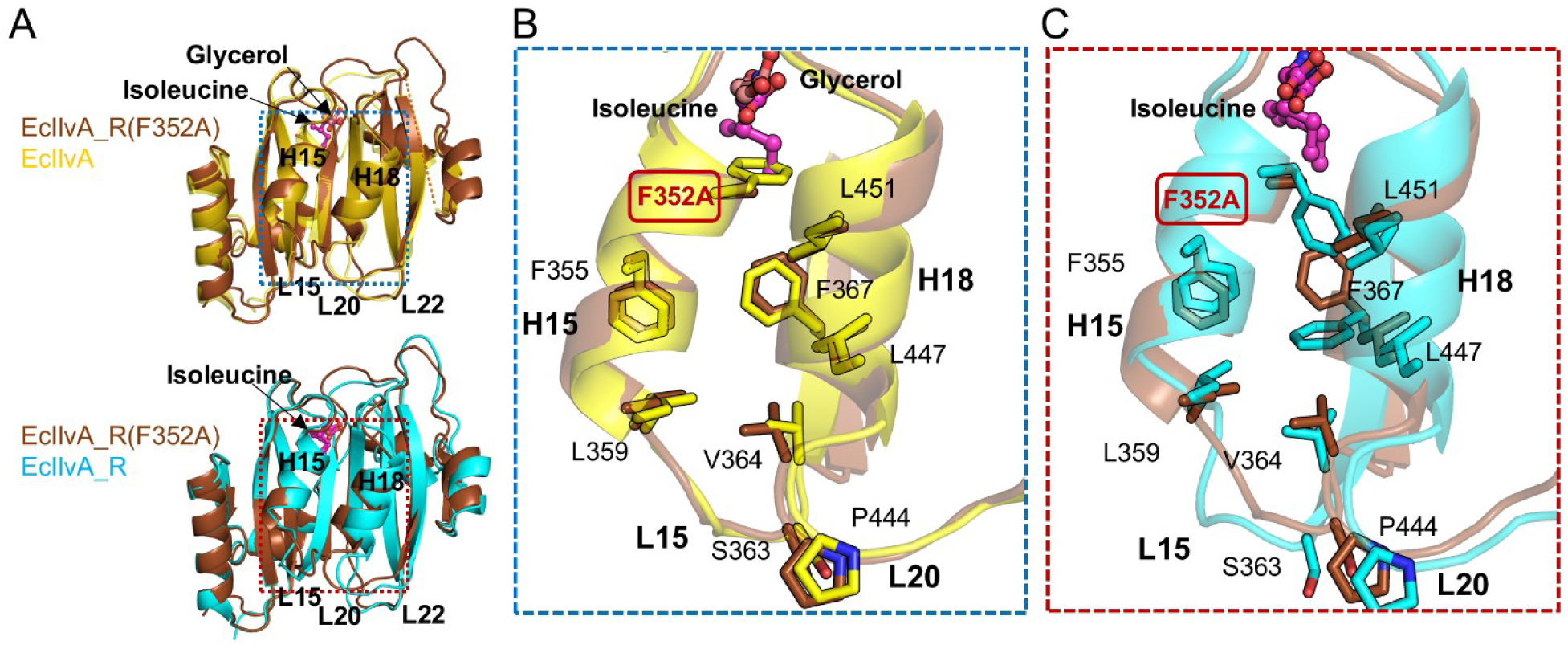
Crystal structure of EcIlvA_R(F352A). *A*, crystal structure of the EcIlvA_R(F352A)•Ile complex (PDB ID: 9D2S). EcIlvA_R(F352A)•Ile is superimposed on EcIlvA (top, PDB ID: 9D2Q) and EcIlvA_R(bottom, PDB ID:9D2R). *B*, close-up view of the isoleucine binding site of EcIlvA_R(F352A)•Ile (brown) and EcIlvA (yellow). *C*, close-up view of the isoleucine binding site in EcIlvA_R(F352A)•Ile (brown) and EcIlvA_R•Ile (cyan).

### Threonine binding at the catalytic site

Although the binding mode of PLP is structurally well characterized in the EcIlvA structures reported here, none contain a bound threonine. Therefore, we docked threonine into the catalytic site, and this identified loop L3, loop L10 and helix H5 as key components to accommodating the substrate (Fig. S6*A*). The amino group of threonine is adjacent to the aldehyde carbon atom of PLP and is hydrogen bonded to the carbonyl oxygen atom of Gly241 in loop L10. The carboxyl group is ideally positioned to interact with the positive dipole at the N-terminus of helix H5, and it also forms hydrogen-bonding interactions with the amide nitrogen atom of His90 and the side chain hydroxyl group of Ser86. Finally, the side chain of threonine interacts with the side chains of Glu240 and Val244 in Loop L10. The location of the active site and the binding of threonine are consistent with allosteric regulation operating across the protomer interface. Most notably, loop L10 directly interacts with loop L20’ in the regulatory domain of the adjacent protomer (Fig. S6*B*). More specifically, the conformational changes of Pro 441’, Ser 443’ and Pro444’ in loop L20’, and D471’ and Y472’ in loop L22’ are well positioned to affect the conformations of Arg234, Leu237, and Phe238 in loop L10 and to impact the binding of the threonine substrate (Figs. S6*B* and S6*C*). We also note that the conformational changes elicited by isoleucine could impact the positioning of the adjacent neck helix H14 and, in turn, the dimer structure (Fig. S6*C*). In terms of the catalytic mechanism, we note that the amino group of the docked threonine is ideally positioned to perform the required initial transamination reaction on the lysine-bound PLP cofactor (*2*) (Fig. S6*A*).

### *Staphylococcus aureus* IlvA - biochemical analysis

Branched-chain amino acids (BCAAs) are important regulators of virulence and fatty acid synthesis in Gram-positive bacteria including *Staphylococcus aureus* (*54*). Inspection of the *S. aureus* IlvA (SaIlvA) sequence revealed that, unlike EcIlvA, there is only one ACT domain in the regulatory domain. It was therefore of interest to investigate the regulation of SaIlvA, if any, by isoleucine. We engineered a SaIlvA full-length (residues 1-422) expression construct (Fig. S1*A*), and the expressed protein was soluble and successfully purified (Fig. S1*D*). The molecular weight based on analytical gel filtration chromatography was estimated to be 380 kDa, which suggests that the SaIlvA assembly is an octamer in solution (Fig. S1*E*). Kinetic analyses of SaIlvA revealed that the regulation of the protein’s activity is completely different from EcIlvA. Isoleucine does not inhibit the binding kinetics of threonine (Fig. 6*A*). The enzyme kinetics data were analyzed using the Michelis-Menten model and the fitted curve closely followed the observed data points, indicating no cooperativity (Fig. 6A). The enzyme also remained active in the presence of 5 mM isoleucine (Figs. 6*A* and 6*B*). In order to analyze and confirm this fundamental difference between SaIlvA and EcIlvA, we used a different assay. Radioactive [^14^C]-threonine was used as a substrate for both proteins and the reaction was resolved by TLC. EcIlvA was inhibited by isoleucine while SaIlvA was not (Fig. S7*A*). The same assay was used to determine if the regulation of SaIlvA involved other intermediates in the branched-chain amino acid pathway. The intermediates and the end product of the isoleucine biosynthesis pathway were tested as inhibitors of SaIlvA and none of the metabolites inhibited SaIlvA activity. These included 2-ketobutyrate (IlvA product), α-keto-β-methylvalerate (the substrate for IlvE), 2-methylbutyrate, 2-methylbutyryl-CoA (product of branched chain keto acid dehydrogenase), isoleucine, valine, serine and anteiso15:0 fatty acid or the end product anteiso15-ACP (Figs. S7*B* and S7*C*). Finally, using the thermal stabilization assay, we confirmed that isoleucine does not bind to SaIlvA (Fig. 6*C*). Unlike EcIlvA, which was increased ∼14 °C in the presence of isoleucine (Fig. *4E*), SaIlvA had nearly identical stabilities to thermal denaturation in both the absence and presence of 5 mM isoleucine (Fig. *6C*). The spectral scans indicate a spectral shift from 410 nm to 430 nm upon threonine binding indicating the external aldimine formation between PLP and threonine for both EcIlvA and SaIlvA (Figs. S7*D* and S7*E*). This shift is similar to other PLP binding enzymes like aspartate aminotransferase that show a shift in the spectra to 430 nm indicating enzyme-substrate interaction (*55, 56*).

**Figure 6.**
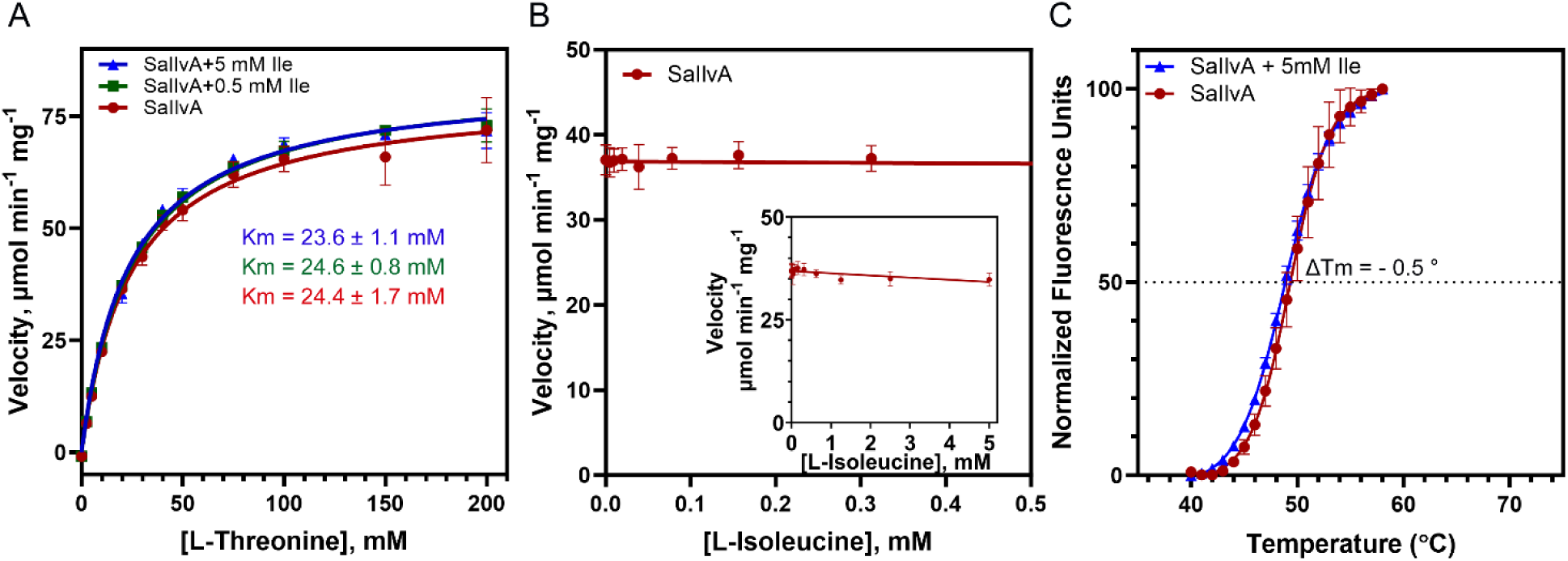
Isoleucine effect on SaIlvA. *A*, effect of isoleucine on the steady-state kinetics of SaIlvA. The theoretical curve was analyzed by the Michaelis-Menten equation using GraphPad Prism 10.1.2. The kcat value was calculated to be 64.4 ± 1.3 sec^-1^. Km and kcat values are presented as mean values ± standard error (SE). *B*, effect of isoleucine on the activity of SaIlvA. *Inset*, plot at the high concentration of isoleucine (0 – 5 mM). The initial velocity was measured in the presence of 30 mM threonine. *C*, the thermal stability of SaIlvA in the absence and presence of 5 mM isoleucine. Data represented as mean ± standard deviation (error bars) from triplicate sets.

### Crystal structure of the *S. aureus* IlvA regulatory domain (SaIlvA_R)

To understand the differences between EcIlvA and SaIlvA, we performed a crystallographic analysis on the latter. Full length SaIlvA failed to generate crystals, but the regulatory domain (SaIlvA_R) was successfully crystallized. The SaIlvA_R gene (residues 336-422) was cloned with an N-terminal 6xHis-tag (Fig. S1*A*) and the expressed protein was soluble and successfully purified (Fig. S1*D*). The protein was estimated to be a 46 kDa tetramer based on gel filtration chromatography (Fig. S1*E*). Crystals were obtained in space group I4_1_, and the structure of SaIlvA_R was determined at 2.8 Å resolution (Table S2). Two tetramers are in the asymmetric unit and are related by a 173° rotation, slightly deviating from perfect 2-fold symmetry, and the RMSD between the two tetramers is 1.2 Å. The tetrameric assembly of SaIlvA_R is consistent with gel filtration analysis and can be considered as a dimer of dimers comprising protomers A/B and C/D (Figs. S1*E* and 7*A*). All eight protomers in the asymmetric unit are structurally very similar with RMSD values between 0.5 to 1.5 Å on Cα atoms in pairwise comparisons.

Unlike EcIlvA_R, which is composed of two ACT domains (ACT1 and ACT2), SaIlvA_R is composed of one ACT domain as predicted from the sequence (Figs. S8*A* and *S8B*) (*57*). Protomers A/B and C/D generate dimers that closely resemble the EcIlvA_R structure (Figs. 7*B* and 7*C*). The overall alignment of the central four sheets (S11, S12, S13, and S14) of the SaIlvA_R monomer was quite good with low RMSD value when overlaid with the isolated ACT1 domain of EcIlvA_R, indicating similarity in their core structures (Fig. 7*D*). The RMSD value for the entire ACT domain is 3.4 Å, whereas for the central four sheets, it is significantly lower at 0.6 Å. SaIlvA_R also has an extra helix (H17) in the prototypical ACT domain, like the ACT1 and ACT2 subdomains of EcIlvA_R (Figs. 7*B* and 7*C*). Sheets S12 and S12′ and helices H15 and H15′ from the partner protomers participate in four key interactions along the dimeric interface: Leu352 interacts with Leu352′, Val356′ and Ile366′; Val356 interacts with Leu352′, Arg353′ and Val356′; Ile366 interacts with Leu352′; Phe369 interacts with Phe369′. In addition, Asn357 and Asn357′ on helices H15 and H15’ form hydrogen bonding interactions (Fig. S9*A*). Protomers A/D and B/C also create dimers, and their interface comprises the additional helices H17 and loops L16 that contain a disordered region (Fig. S9*B*).

**Figure 7.**
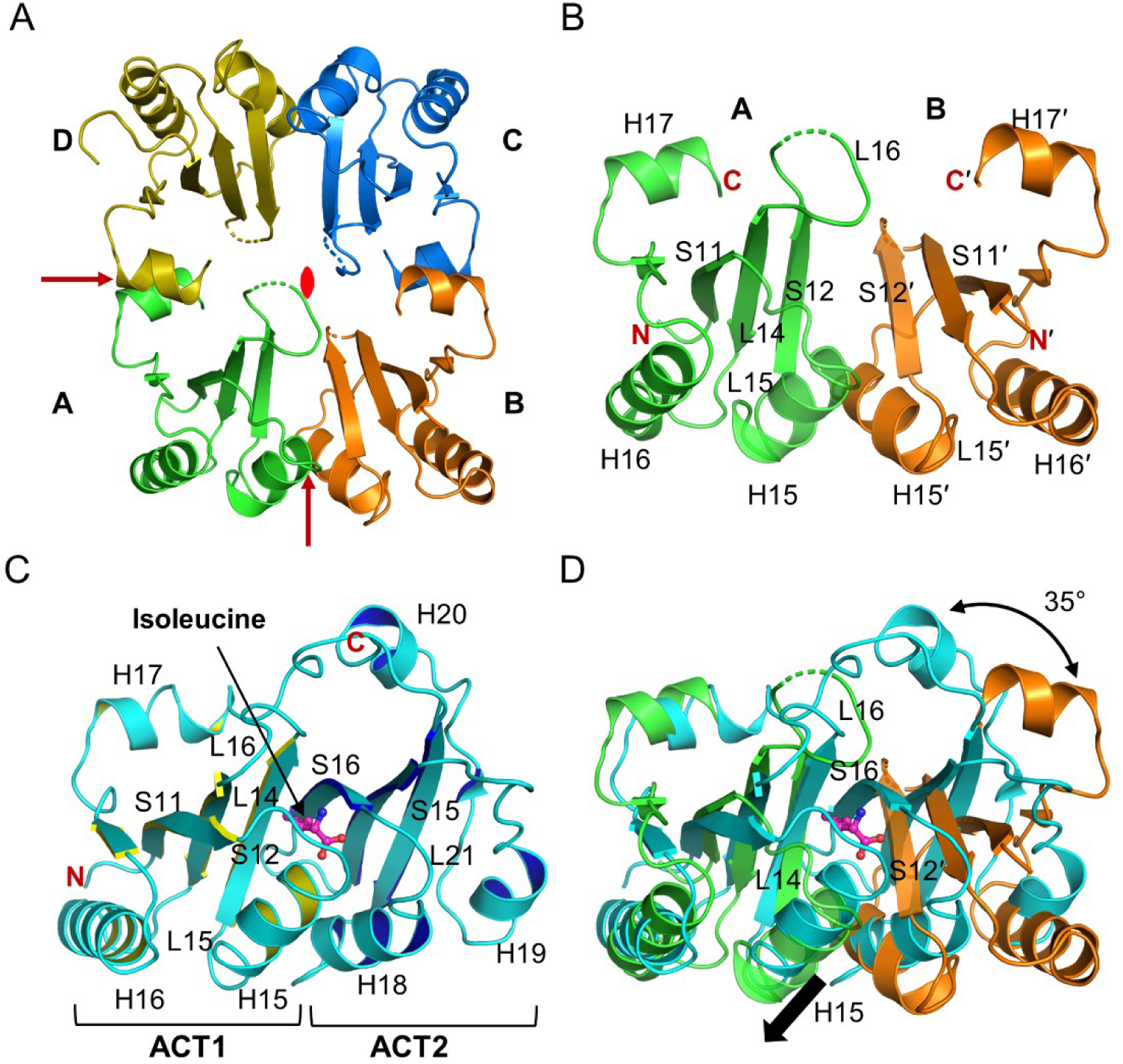
Crystal structure of SaIlvA_R. *A*, tetramer of the SaIlvA_R crystal structure (PDB ID: 9D2T). Each monomer of the tetramer is related by an approximate twofold axis (read symbol and arrows). Two tetramers are in the asymmetric unit. *B*, dimeric structure of SaIlvA_R. Protomers A and B are shown in green and orange, respectively. *C*, monomeric structure of EcIlvA_R•Ile (PDB ID: 9D2R). The two ACT domains are shown in cyan ribbon with highlighted yellow and blue. The bound isoleucine is shown with magenta balls-sticks. *D*, the overlayed structures of dimeric SaIlvA_R (green and orange) and monomeric EcIlvA_R•Ile (cyan). The protomer A of SaIlvA_R is superimposed on the ACT1 domain of EcIlvA_R•Ile. The conformational difference of helix H15 is shown as a black arrow.

Despite the structural similarities between EcIlvA_R and the SaIlvA_R A/B and C/D dimers, key differences appear to prevent isoleucine binding to the latter. Helix H15 and loops L14, L15 and L16 exhibit different conformations compared to their EcIlvA_R counterparts, and the H15 and L14 counterparts play crucial roles in the binding of isoleucine in EcIlvA (Figs. 7*B* and 7*C*). Also, the relative orientations of the ACT1 and ACT2 subdomains of EcIlvA_R and the A/B and C/D protomers of SaIlvA_R are significantly different; when ACT1 of EcIlvA_R is aligned with protomer A, protomer B is rotated ∼35° from the ACT2 domain of EcIlvA_R (Fig. 7*D*). Isoleucine binds at the interface of ACT1 and ACT2 in EcIlvA_R and the different orientations in the A/B and C/D dimers of SaIlvA_R disrupt the isoleucine binding site (Fig. 7*D*). More specifically, the different location of sheet S12′ in protomer B corresponding to sheet S16 of the ACT2 domain of EcIlvA disrupts the binding of isoleucine in SaIlvA (Fig. 7*D*). The conformation of helix H15 in SaIlvA_R is different from that in EcIlvA_R, so the isoleucine model obtained by overlaying the entire an ACT domain could not interact with helix H15 (Fig. 7D). Therefore, even when SaIlvA_R and EcIlvA_R•Ile were overlaid only on helix H15 to locate the isoleucine that could interact with helix H15 in SaIlvA_R, the binding of isoleucine on SaIlvA was not favorable (Figs. S9*C* and S9*D*). Specifically, Glu347 interacts with the amino group of the isoleucine in the EcIlvA_R complex, whereas the corresponding residue, Gln347, in the SaIlvA_R is oriented toward the solvent. Two other isoleucine interacting residues, Asn459 and Ile460, are located in sheet S16 of EcIlvA_R, whereas their counterparts, Asp365′ and Ile366′, are located in sheet S12′, which has a different orientation compared to sheet S16 in EcIlvA_R. Leu352 in SaIlvA_R corresponds to Phe352 in EcIlvA, a key residue for the feedback inhibition by isoleucine (Figs. S9*C* and S9*D*). This analysis indicates that isoleucine binding is not favorable in SaIlvA and therefore unregulated by isoleucine.

### Structural model of the SaIlvA assembly

In the absence of a full-length SaIlvA crystal structure, we predicted the assembly using a combination of the protomer AlphaFold model (AF-Q2FF63) and our crystal structure of the SaIlvA_R tetramer. The AlphaFold model was first compared with the crystal structure of EcIlvA (PDB ID: 9D2Q) and the catalytic domains are very similar, consistent with their 36.3% sequence identity (Figs. S8*A* and S8*B*). The structure of the regulatory domain of the model is also very similar to the domain observed in the crystal structure of SaIlvA_R (PDB ID: 9D2T), except for loops L14 and L16 (Fig. S8*C*). Thus, the individual domains of AlphaFold model appear to be excellent. However, when the model is superimposed on the tetrameric SaIlvA_R to generate the full assembly, there is no close contact between the catalytic domains (Fig. S10*A*). Two domains of SaIlvA are connected by a linker, allowing for potential flexibility in their relative orientation. To obtain an improved oligomeric structure, the AlphaFold model of the dimeric SaIlvA was first predicted using ChimeraX and ColabFold (Fig. S10*B*) (*48, 58*). The catalytic domain assembly of the resulting dimeric SaIlvA model is similar to that of the dimeric EcIlvA with interactions between the catalytic domains (Figs. S10*B* and S10*C*). In the crystal structure of SaIlvA_R, the N-termini of protomers A and C align in the same direction, while those of the protomers B and D face the opposite direction (Fig. 8*A*). Therefore, when the model of the dimeric SaIlvA was superimposed on protomers A/C and B/D of SaIlvA_R (Fig. 8*B*), the resulting tetramer model has a central regulatory domain assembly with paired catalytic domains extending in opposite directions (Fig. 8*C*). This is opposite of the EcIlvA assembly in which the catalytic domains constitute the core of the tetramer (Figs. 2*B* and 8*C*). Due to the agreement between copies in the asymmetric unit, it is unlikely that this organization is due to a crystallographic artifact.

**Figure 8.**
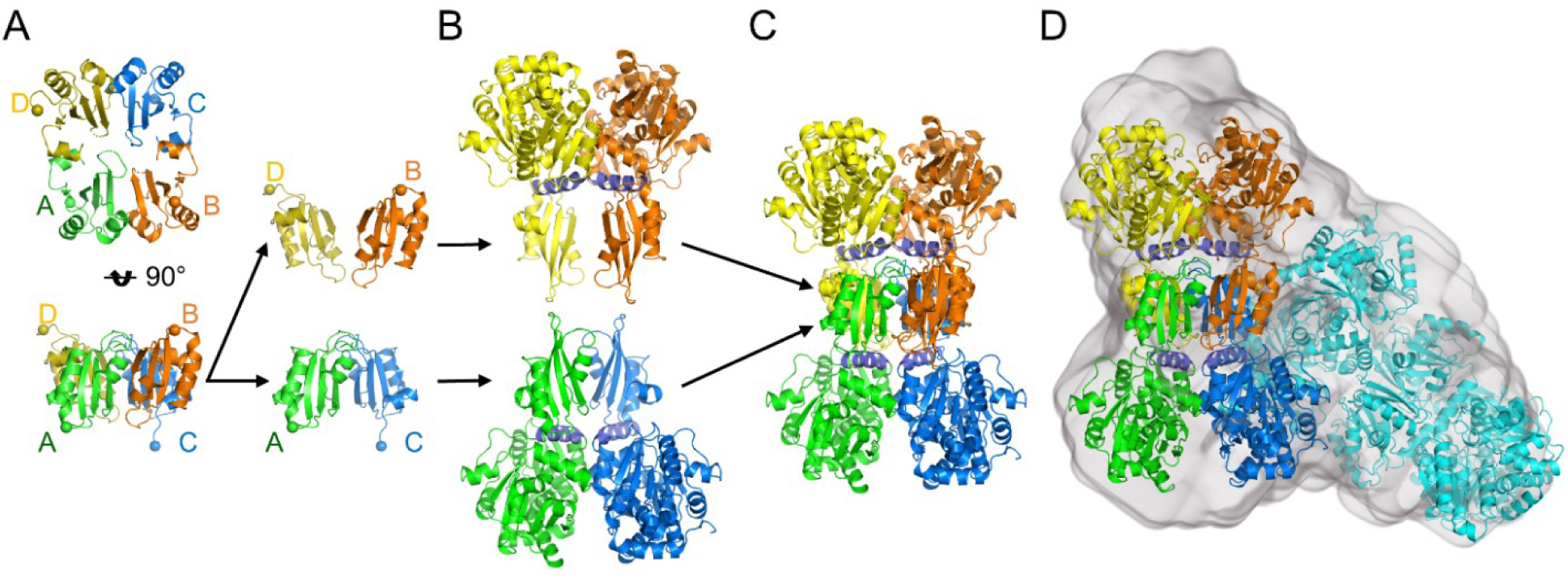
Modelled quaternary structure of SaIlvA. *A*, tetramer of SaIlvA_R is shown in green (protomer A), orange (protomer B), blue (protomer C), and yellow (protomer D). The N-terminus of each protomer is shown as a sphere. The N-termini of protomers A and C align in the same direction, while those of the protomers B and D face the opposite way. *B*, dimer of SaIlvA generated using AlphaFold. The regulatory domains of SaIlvA dimer can be overlayed on protomers A/C and B/D. *C*, SaIlvA tetramer model was generated using AlphaFold model of SaIlvA dimer and SaIlvA_R tetramer crystal structure. The four C-terminal regulatory domains form the core of the tetramer. *D*, the SaIlvA octamer model, which consists of two tetramers, fitted into the electron density determined by small angle X-ray scattering (SAXS) in solution. A tetramer model is shown in green, orange, blue, and yellow, and the other tetramer is shown in cyan.

As noted above, gel filtration of the purified SaIlvA suggested an octameric assembly but the modeling suggests a tetramer. To resolve this issue, we used SEC-SAXS analysis to determine the volume occupied by SaIlvA in solution (Figs. 8*D*, S4*C*, S4*F*, S4*I*, and Table S3). Comparison of the SaIlvA SAXS experimental data set with the predicted profile of the generated tetrameric model of SaIlvA is very poor, but the SAXS data fit well to the theoretical behavior of a SaIlvA octamer (χ^2^ = 1.37) (Fig. S4*I*). Specifically, the SAXS analysis is consistent with SaIlvA existing as an octamer with a molecular weight of 321 kDa in solution, which agrees well with the gel filtration profile (Table S3 and Fig. S1*E*). A SaIlvA octamer, consisting of two associated tetramers, was constructed that optimally fits the *ab initio* electron density maps generated from the solution scattering data, with the correlation coefficient of 0.79 (Fig. 8*D*).

### Complementation of a SaIlvA knockout strain by EcIlvA and EcIlvA(F352A)

The Δ*SaIlvA* knockout strain PDJ78 was transformed by three plasmids, pCN38 (empty vector) (*28*), pCN38 SaIlvA or pCN38 EcIlvA. All the three strains grew in ‘complete defined medium’ (CDM) that contains isoleucine as expected, and the strain complemented with pCN38 SaIlvA also grew in CDM minus isoleucine consistent with the presence of the active enzyme that lacks feedback inhibition. However, due to the lack of IlvA enzyme the strain complemented with pCN38 could not grow in the absence of isoleucine. The strain complemented with pCN38 EcIlvA also could not grow well in the absence of isoleucine as it got feedback regulated by the newly synthesized isoleucine and hence could not keep up with the demand for isoleucine needed for membrane synthesis. (Fig. 9*A*). To understand the effect of the feedback regulation on fatty acid synthesis in these three strains, the cells at the end of the growth curve shown in panel A were harvested and their phospholipids were extracted. The fatty acids from the phospholipids were transformed into methyl esters and subjected to gas chromatography. In CDM, the fatty acid composition was remarkably indistinguishable in the three strains with a slight increase in straight chain odd fatty acids in the strain with SaIlvA (Fig. 9*B*). However, in the absence of isoleucine, there was an increase in *anteiso* odd chain fatty acids in the pCN38 SaIlvA strain compared to the strain with pCN38 EcIlvA. This indicates that isoleucine was copiously synthesized to engage in fatty acid synthesis with SaIlvA present but not with EcIlvA, again confirming feedback inhibition with the latter (Fig. 9*C*). Also, an increase in straight chain even fatty acid was observed with the EcIlvA indicating that the decrease in *anteiso* odd chain fatty acids was compensated with an increase in straight chain even fatty acids in the absence of isoleucine (Fig. 9*C*). This change in the *anteiso* to straight chain fatty acids would affect the membrane fluidity preventing the growth of *S. aureus* when complemented with EcIlvA.

**Figure 9.**
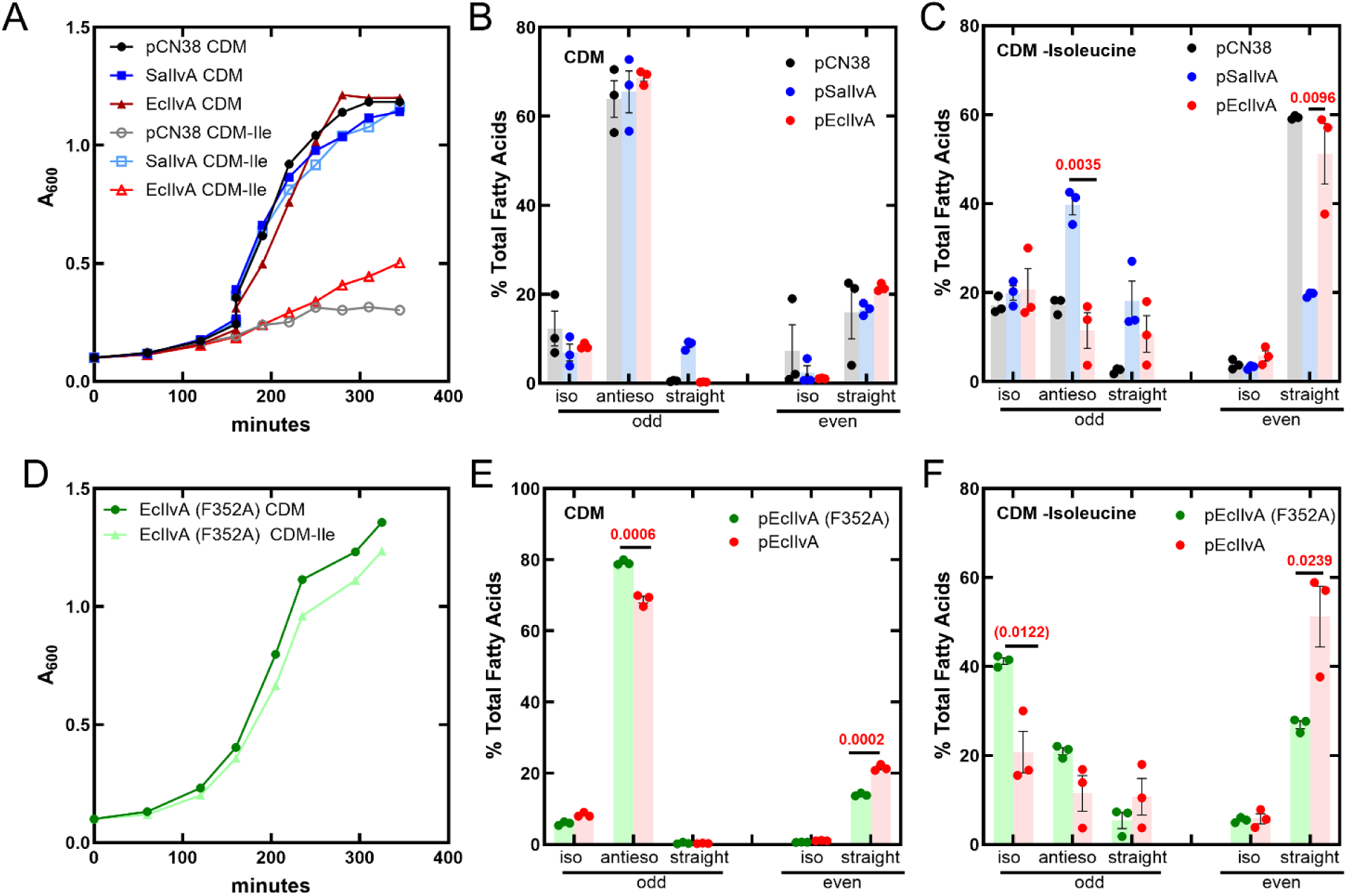
Complementation of Δ*IlvA*. *A*, growth curve of Δ*IlvA* containing empty vector (pCN38) or pSaIlvA or pEcIlvA grown in Complete Defined Medium (CDM) or CDM minus Isoleucine. *B*, fatty acid analysis by Gas Chromatography of Δ*IlvA* containing empty vector (pCN38) or pSaIlvA or pEcIlvA grown in CDM. *C*, fatty acid analysis by Gas Chromatography of Δ*IlvA* containing empty vector (pCN38) or pSaIlvA or pEcIlvA grown in CDM minus Isoleucine. *D*, growth curve of Δ*IlvA* containing pEcIlvA (F352A) mutant grown in Complete Defined Medium (CDM) or CDM minus Isoleucine. *E*, fatty acid analysis by Gas Chromatography of Δ*IlvA* pEcIlvA (F352A) mutant grown in CDM. *F*, fatty acid analysis by Gas Chromatography of Δ*IlvA* pEcIlvA (F352A) mutant grown in CDM minus Isoleucine. The data in *A* and *D* are representative of experiments done twice. The data in *B*, *C*, *E*, and *F* are from triplicate sets (mean ± SE). The value for the t-test is given in red.

To further analyze the regulation of isoleucine synthesis by EcIlvA, we expressed the F352A mutant of EcIlvA (pCN38 EcIlvA (F352A)) in the Δ*SaIlvA* knockout. The growth curves of this strain clearly show that it can grow in both the presence and absence of isoleucine (Fig. 9*D*), which is not the case with the EcIlvA strain (Fig. 9*A*). The fatty acid analysis (Figs. 9*E* and 9*F*) shows that there is a significant increase in *anteiso* odd chain fatty acids and a decrease in the straight chain even fatty acids in the EcIlvA (F352A) compared to the EcIlvA even when grown in CDM. A significant decrease in the straight chain even fatty acids was observed when grown in the absence of isoleucine. The branched-chain fatty acids made due to the lack of feedback regulation in the EcIlvA (F352A) mutant allow the Δ*SaIlvA* knockout to grow in the absence of isoleucine in the media. These results confirm the importance of F352 in regulating isoleucine synthesis by EcIlvA.

## DISCUSSION

The majority of Gram-positive bacteria, including *S. aureus*, incorporate branched-chain *iso* and *anteiso* fatty acids into their membrane as crucial components that contribute to and control membrane fluidity (*59, 60*). More specifically, isoleucine derived exogenously or synthesized *de novo* is converted to keto acids and diverted into FASII. It has been shown that removal of isoleucine from the media increases *iso* fatty acids to maintain membrane fluidity (*9*). This key role of isoleucine in *S. aureus* prompted us to analyze SaIlvA and its regulation. IlvA from *E. coli* (EcIlvA) has been extensively studied and was the first enzyme identified that displays allosteric control whereby isoleucine modulates its activity via a binding site distinct from the active site and thereby controls throughput of the BCAA pathway. Structural analysis of EcIlvA has revealed a dimer-dimer tetrameric assembly in which each protomer comprises two domains, and biochemical analyses revealed that one domain regulates the activity of the catalytic site that resides in the second domain. The sequence of SaIlvA suggests that the regulatory domain has a different architecture from that of EcIlvA, and the overall goal of this study was to investigate the structure, biochemistry and regulation of SaIlvA and to establish whether it also allosterically controls the BCAA pathway in this organism and in Gram-positive bacteria generally.

Despite the wealth of knowledge surrounding the biochemistry and structure of EcIlvA, neither the isoleucine binding site within the IlvA regulatory domain nor the structural consequences of isoleucine binding have been directly observed. Our studies therefore initially focused on EcIlvA. Although we failed to structurally characterize full-length EcIlvA in its isoleucine-bound inactive state, we succeeded in determining the crystal structure of the EcIlvA_R regulatory domain bound to isoleucine and identified conformational changes that are propagated across the structure via a classic allosteric pathway. The direction of the pathway strongly suggests that the allosteric regulation occurs across the dimer interface with the isoleucine binding to the EcIlvA_R of one protomer negatively regulating the active site of the catalytic module of the adjacent protomer (Fig. 10). The structural data that support this are twofold. First, the active site in the catalytic module is 43 Å away from the isoleucine site on the partner protomer regulatory domain but 54 Å away from its own regulatory domain (Fig. 10). Second, the isoleucine binding signal is transmitted 23 Å from the isoleucine binding site to the L20 loop, which is located at the dimer interface and directly contacts the partner active site (Figs. 10 and S6*B*). This proposal that EcIlvA allostery operates within a quaternary structure that undergoes small but significant conformational changes upon effector binding is a basic tenet of the fundamental theory. Although we were not able to structurally characterize these presumed changes, they may involve the conformation of the neck helix H14 that connects the regulatory and catalytic subdomains of EcIlvA (Figs. 10 and S6*C*) and is in a position to be affected by the allosteric pathway. When the previously determined crystal structure of EcIlvA is compared with the higher resolution structure presented here, the catalytic and regulatory domains flanking the neck helix are in slightly different relative positions which suggests a degree of interdomain flexibility within the protomer. Dynamic control of the neck helix conformation and the quaternary structure upon isoleucine binding may explain why it was not possible to crystallize the inactive isoleucine-bound EcIlvA.

**Figure 10.**
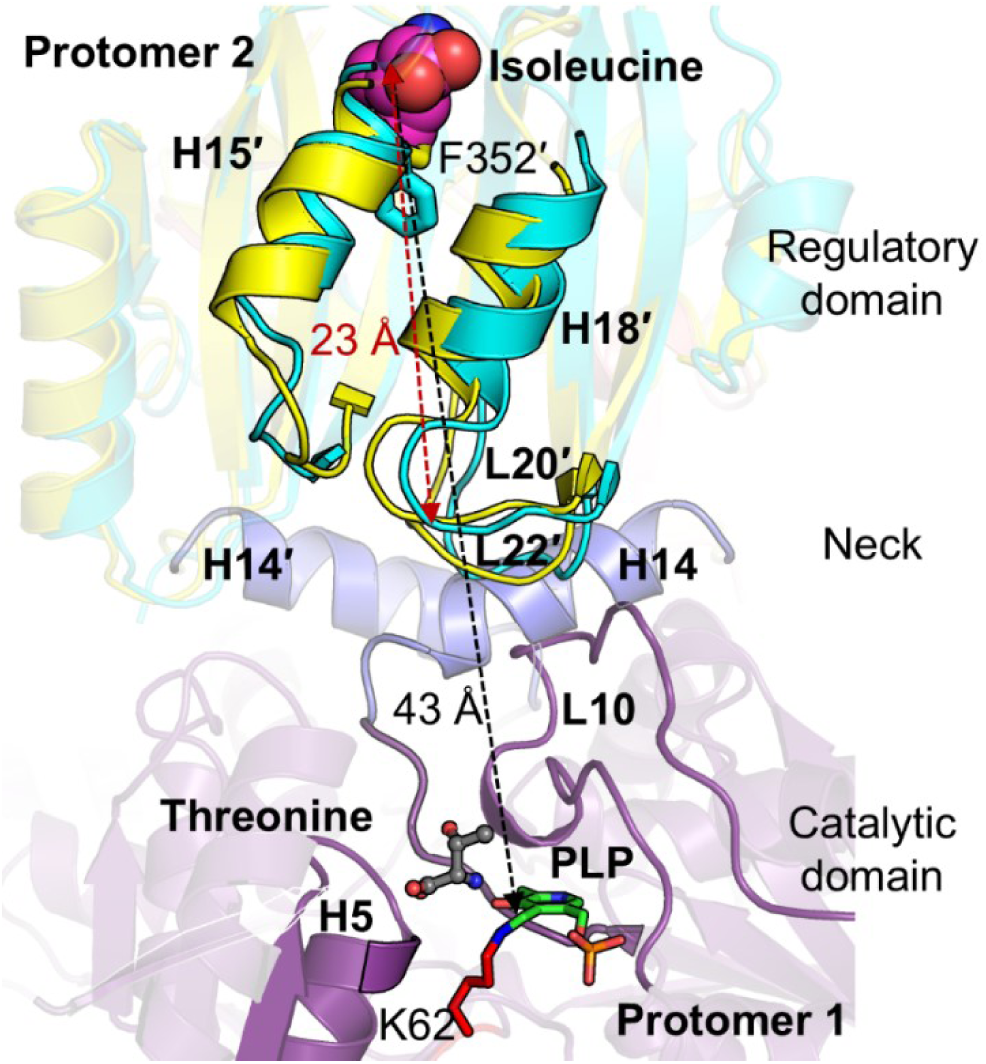
Allosteric regulation of isoleucine on EcIlvA. Loop interactions between the catalytic and regulatory domain in the active site of EcIlvA. The catalytic domain of protomer 1, regulatory domain of protomer 2 and necks of both protomers of EcIlvA are shown in purple, yellow, and blue, respectively. The EcIlvA_R•Ile structure (cyan cartoon) is superimposed on the EcIlvA structure. The features of protomer 2 are indicated with a prime. The docked threonine molecule is represented by ball-and-sticks with gray carbon atoms, blue nitrogen atom, and red oxygen atoms. PLP and the active-site lysine (K62) are shown as sticks. The red arrow indicates the distance from isoleucine to L20 loop, while the black arrow indicates the distance from isoleucine to PLP of the partner protomer.

The identification of Phe352 as the key residue at the start of the allosteric pathway in EcIlvA_R directly adjacent to the bound isoleucine is an important finding that we tested by biochemically and structurally characterizing the F352A point mutant (Figs. 4, 5 and 10). These analyses confirmed the importance of this residue. EcIlvA(F352A) still binds isoleucine, but the structure of EcIlvA_R(F352A) is very similar to EcIlvA_R and the allosteric pathway is unaffected. Thus, F352A prevents the initial communication as predicted. Also, EcIlvA(F352A) became insensitive to inhibition by isoleucine despite clearly binding the effector. A number of studies have identified point mutants that render EcIlvA refractory to inhibition by isoleucine (*19, 20, 25, 61*). These residues are shown in Fig. S11 and they all occur within the regulatory domain, either directly affecting the isoleucine binding site or the allosteric pathway. It was reported that the ligand binding site mutants E347F, F352A, and I460F became remarkably insensitive to isoleucine inhibition, with increased IC50 values by factors 43, 35, and 49, respectively (*19*). Arg362 in loop L15 makes a salt bridge with Glu394 in the EcIlvA_R structure. The R362F mutant, lacking salt bridge, became strongly feedback-resistant with an increased IC50 value of 104-fold compared to that of the wild-type EcIlvA (*19*). Furthermore, previous studies with EcIlvA, mutations in residues P441L, G445E, A446E, L447A, L451A, and L454A, which are in the allosteric pathway, showed resistance to isoleucine inhibition (*19, 20, 25, 61*). It should be noted that the location of the isoleucine binding site is different from the binding site that was inferred by computational modeling and mutagenesis (*20*).

Having a much clearer understanding of how allostery operates in EcIlvA, we then studied the process in SaIlvA. We clearly demonstrated that SaIlvA is not allosterically controlled by isoleucine and indeed does not bind isoleucine. The possibility that SaIlvA and the associated BCAA pathway are controlled by another metabolite, possibly associated with phospholipid biosynthesis, was also tested and none were identified. Our studies therefore strongly suggest that allosteric feedback inhibition is not present in SaIlvA (Fig. 9), and the essentiality of isoleucine to membrane phospholipids in *S. aureus* may explain why. We failed to crystallize full-length SaIlvA but succeeded in determining the crystal structure of SaIlvA_R. It comprises a tetrameric assembly with two dimers that each resemble the EcIlvA_R protomer, but the isoleucine binding site is not present due to differing orientations of the paired ACT domains. The tetrameric assembly of the ACT domains in SaIlvA_R reflects the variety of ways in which this domain can generate multimers (*18, 62, 63*). Modeling of the full-length SaIlvA structure suggests that the tetrameric assembly of SaIlvA_R forms the core of the SaIlvA tetrameric assembly, unlike EcIlvA where the catalytic domain creates the core of the tetramer. This further suggests that it is the catalytic domains that extend outwards from the regulatory domains, which is opposite from the EcIlvA assembly. SEC-SAXS data and gel filtration profiles both agree that SaIlvA exists as an octamer in solution. The SEC-SAXS analyses are consistent with two associated tetrameric assemblies, but we have no clear model to explain this. Although the crystal structure of SaIlvA_R has two tetramers in the crystal asymmetric unit, the optimal SaIlvA octamer based on the SEC-SAXS is not consistent with this asymmetric unit, which is therefore unlikely to structurally determine the SaIlvA octamer. Finally, complementation studies of Δ*SaIlvA* in which IlvA has been removed confirmed our conclusions based on the biochemical and structural analyses of SaIlvA. Notably, Δ*SaIlvA* complemented with EcIlvA grew very poorly, consistent with the idea that allosteric control of IlvA severely limits the growth of *S. aureus* via reduced membrane synthesis due to a deficiency in essential anteiso acyl chains and hence its absence from SaIlvA.

IlvA plays a crucial role not only in BCAA production in bacteria but also in plants, offering commercial opportunities for amino acid overproduction through fermentation (*64, 65*) and for enhancing crop production (*66, 67*). Isoleucine, an essential amino acid for humans, is vital for immunity, blood sugar regulation, and hormone production (*68–71*). Our analysis of IlvA’s feedback inhibition mechanism and the key residues involved in its active and allosteric binding sites opens new possibilities for developing small molecule effectors and engineered variants to enhance isoleucine production. For instance, our experiments with the EcIlvA (F352A) mutant demonstrated that it can bypass isoleucine feedback regulation, leading to increased isoleucine biosynthesis during fermentation. Additionally, we showed in *S. aureus* that combining IlvA sequence analysis from other bacteria with AlphaFold structural predictions provides valuable insights into their allosteric regulation. This approach could potentially be extended beyond bacteria to crops, enhancing their nutritional and therapeutic value (*72*).

## DATA AVAILABILITY

The atomic coordinates and structure factors have been deposited in the worldwide Protein Data Bank (EcIlvA, PDB ID: 9D2Q; EcIlvA_R•Ile, PDB ID: 9D2R; EcIlvA_R(F352A)•Ile, PDB ID: 9D2S; and SaIlvA_R, PDB ID: 9D2T). SAXS data and models have been deposited in the Small Angle Scattering Biological Data Bank (SASDBD, https://www.sasbdb.org/) (*73*) under the accession codes SASDVC7 (EcIlvA), SASDVD7 (EcIlvA_R), and SASDVE7 (SaIlvA). All other data produced for this work are contained within the article.

## ACCESSION CODES

EcIlvA: P04968 SaIlvA: Q2FF63

## SUPPORTING INFORMATION

List of primers and plasmids (Table S1); data collection and refinement statistics for crystal structures (Table S2); macromolecular properties of IlvA and IlvA_R from SEC-SAXS (Table S3); design and purification of IlvA protein constructs (Figure S1); structural overview of EcIlvA (Figure S2); improved model in three regions of EcIlvA structure (Figure S3); analysis of the IlvA solution dynamics by SEC-SAXS (Figure S4); electron density maps (Figure S5); threonine binding at the catalytic site (Figure S6); EcIlvA and SaIlvA assay with threonine (Figure S7); structural overview of SaIlvA_R (Figure S8); the dimeric structure of SaIlvA_R (Figure S9); AlphaFold model of SaIlvA (Figure S10); structural location of mutations reported to render EcIlvA refractory to isoleucine inhibition (Figure S11) (PDF).

## FUNDING AND ADDITIONAL INFORMATION

This work was supported by National Institutes of Health grants AI166116 (C.D.R.), GM034496 (C.O.R.), Cancer Center Support Grant CA21765, and ALSAC, St. Jude Children’s Research Hospital. The content is solely the responsibility of the authors and does not necessarily represent the official views of the National Institutes of Health.

## ACKNOWLEDGMENTS

We thank the St. Jude Children’s Research Hospital (SJCRH) Biomolecular X-Ray Crystallography Center for support with X-ray data collection, and the SJCRH Hartwell Center DNA Sequencing Shared Resource for DNA sequencing. We particularly thank Dr. Stephen W. White for thoughtful suggestions and valuable input in developing this manuscript.

The X-ray data were collected at three shared resources. (1) The Advanced Photon Source (APS) is a U. S. Department of Energy (DOE) Office of Science user facility operated for the DOE Office of Science by Argonne National Laboratory under Contract No. DE-AC02-06CH11357. Diffraction data were collected at the Southeast Regional Collaborative Access Team (SER-CAT) 22-ID beamline. SER-CAT is supported by its member institutions (https://www3.ser.aps.anl.gov/contact-us#TITLE_SER_CAT_Memberships), equipment grants (S10_RR25528, S10_RR028976 and S10_OD027000) from the National Institutes of Health, and funding from the Georgia Research Alliance. SAXS data were collected at BioCAT (18-ID beamline), which is supported by grant P30 GM138395 from the National Institute of General Medical Sciences of the National Institutes of Health. (2) Diffraction data were collected at the Center for BioMolecular Structure (CBMS) at the National Synchrotron Light Source II at Brookhaven National Laboratory and is primarily supported by the National Institutes of Health, National Institute of General Medical Sciences (NIGMS) through a Center Core P30 Grant (P30GM133893), and by the DOE Office of Biological and Environmental Research (KP1607011). The National Synchrotron Light Source II is operated under Contract No. DE-SC0012704 for the U.S. Department of Energy (DOE), Office of Science. (3) Diffraction data were collected at beamline 5.0.2 of the Advanced Light Source, a DOE Office of Science User Facility at Lawrence Berkeley Laboratory under Contract No. DE-AC02-05CH11231, and is supported in part by the ALS-ENABLE program funded by the National Institutes of Health, National Institute of General Medical Sciences, grant P30 GM124169-01.

## ABBREVIATIONS

### The abbreviations used are

IlvA: threonine deaminase
BCAA: branched-chain amino acids
PLP: pyridoxal 5′-phosphate
PGDH: phosphoglycerate dehydrogenase
ACT domain: a structural motif of 70-80 amino acids.

