## Supporting Information for "Isoleucine binding and regulation of *Escherichia coli* and *Staphylococcus aureus* threonine dehydratase (IlvA)"

**This PDF files includes:**

SI Tables S1-S3

SI Figures S1-S11

**Table S1. List of primers and plasmids**

| Primer | Sequence |
|---|---|
| EcIlvA-NdeI-F | GCCGCGCGGCAGCCATGCTGACTCGCAACCC |
| EcIlvA-EcoRI-R | CTTGTCGACGGAGCTCGAATTCCTAACCCGCCAAAAAGAACC |
| EcIlvA-reg-NdeI-F | GCCGCGCGGCAGCCATGAACAGCGTGAAGCGTTG |
| EcIlvA-reg-EcoRI-R | CTTGTCGACGGAGCTCGAATTCCTAACCCGCCAAAAAGAACC |
| EcIlvA(F352A)-F | TCCGGAAGAAAAAGGCAGCGCCCTCAAATTCTGCCAACTG |
| EcIlvA(F352A)-R | CAGTTGGCAGAATTTGAGGGCGCTGCCTTTTTCTTCCGGA |
| SaIlvA-NcoI-F | CTTTAAGAAGGAGATATACCATGACAGTCAAAACAACAGTTT |
| SaIlvA-XhoI-R | TGGTGGTGGTGGTGCTCGAGAATTAACAATGAATATAACATCTTATTTTC ATTAATA |
| SaIlA-reg-NdeI-F | GCCGCGCGGCAGCCATGAAATGAAGCATTACTTTATCTTAAATTTCC |
| SaIlvA-reg-EcoRI-R | CTTGTCGACGGAGCTCGAATTCGACTGTTTTTCTTACTATGTGTTAAAT |
| SaIlvA-UP-EcoRI-F | GCCCTTTCGTCTTCAAGAATTCTTAGGGTATGTTTTTCTCGGTTC |
| SaIlvA-UP-R | ACTATGTGTTACTGTCATAATATTCAACTCCC |
| SaIlvA-DN-F | TTATGACAGTAACACATAGTAAGAAAAACAGTC |
| SaIlvA-DN-KpnI-R | TCTAGAGGATCCCCGGGTACCGACTTAGTGATCCGGTGG |
| pCN38-SphI-SarAP1-F | AGCTGGCGGCCGCTGCATGCAACAGCTATGACATGATTACGA |
| SarAP1-EcIlvA-R | TGCCCATGGATCCACCTCCTTTCTAG |
| SarAP1-EcIlvA-F | GGTGGATCCATGGGCAGCAGC |
| EcIlvA-pCN38-R | AGCTCGGTACCCGGGGATCTTCCTAACCCGCCAAAAAG |
| SarAP1-SaIlvA-R | GACTGTCATGGATCCACCTCCTTTCTAG |
| SarAP1-SaIlvA-F | GGTGGATCCATGACAGTCAAAACAACAG |
| SaIlvA-pCN38-R | AGCTCGGTACCCGGGGATCTTTCGGGCTTTGTTAGCAG |

| Plasmid Name | Description |
|---|---|
| pPJ669 | pET28a containing full-length *IlvA* gene from E. coli with N-terminal His tag |
| pPJ668 | pET28a containing the regulatory domain of *IlvA* gene from E. coli (amino acid 335-514) with N-terminal His-tag |
| pPJ675 | pET28a-IlvA (F352A) mutant in full-length *IlvA* gene from E. Coli with N-terminal His tag |
| pKM356 | pET28a-IlvA-reg (F352A) mutant in the regulatory domain of *IlvA* gene from E. coli |
| pPJ665 | pET28a containing full-length *IlvA* gene from SaUSA300 |
| pPJ666 | pET28a containing the regulatory domain of *IlvA* gene from SaUSA300 (amino acid 336-422) |
| pPJ676 | allelic replacement plasmid for generating *SaIlvA* knockout in AH1263 |
| pCN38 | Shuttle vector |
| pPJ681 | pCN38 containing full-length His-*EcIlvA* |
| pPJ680 | pCN38 containing full-length *SaIlvA*-His |

**Table S2. Data collection and refinement statistics for crystal structures**

| Parameter | EcIlvA | EcIlvA_R•Ile | EcIlvA_R (F352A)•Ile | SaIlvA_R |
|---|---|---|---|---|
| **Data Collection** [a] | | | | |
| Wavelength (Å) | 1.0 | 0.920 | 1.0 | 1.0 |
| Space group | I222 | $P3_221$ | C2 | $I4_1$ |
| Cell dimensions | | | | |
| a (Å) | 80.0 | 99.4 | 106.4 | 88.9 |
| b (Å) | 93.1 | 99.4 | 39.1 | 88.9 |
| c (Å) | 161.5 | 50.1 | 56.6 | 213.2 |
| α (°) | 90.0 | 90.0 | 90.0 | 90.0 |
| β (°) | 90.0 | 90.0 | 117.0 | 90.0 |
| γ (°) | 90.0 | 120.0 | 90.0 | 90.0 |
| Resolution range (Å) | 60.7-1.87 (1.90-1.87) | 28.7-1.60 (1.64-1.60) | 47.4-1.22 (1.24-1.22) | 44.5-2.80 (2.87-2.80) |
| $R_{merge}$ | 0.127 (3.440) | 0.050 (0.628) | 0.031 (0.195) | 0.084 (0.730) |
| CC1/2 | 0.997 (0.719) | 1.000 (0.787) | 1.000 (0.984) | 0.997 (0.790) |
| Completeness (%) | 99.7 (94.2) | 99.7 (96.5) | 97.9 (89.5) | 99.9 (98.7) |
| Redundancy | 13.4 (13.3) | 8.2 (5.3) | 6.5 (5.3) | 13.5 (9.8) |
| $I/\sigma(I)$ | 10.0 (1.1) | 20.9 (2.0) | 28.4 (7.1) | 19.0 (2.1) |
| Unique reflections | 50,171(2,340) | 37.700 (2,684) | 60,600 (2,718) | 20,172 (1,410) |
| **Refinement** | | | | |
| Resolution range (Å) | 60.7-1.87 | 28.7-1.60 | 47.4-1.22 | 44.5-2.805 |
| No. of reflections | 50,144 | 37,679 | 60,587 | 20,118 |
| No. of atoms | | | | |
| Protein | 3,976 | 1,453 | 1,459 | 5,415 |
| Ligand | 58 | 43 | 15 | |
| Water | 228 | 198 | 149 | 14 |
| $R_{work}$ | 0.176 | 0.153 | 0.141 | 0.192 |
| $R_{free}$ | 0.202 | 0.166 | 0.151 | 0.239 |
| Average B factors | | | | |
| Protein | 46.2 | 27.5 | 19.8 | 93.3 |
| Ligand | 60.5 | 37.6 | 18.1 | |
| Water | 51.0 | 41.1 | 31.0 | 65.1 |
| Rmsd from ideal | | | | |
| Bond lengths (Å) | 0.006 | 0.013 | 0.011 | 0.002 |
| Bond angles (˚) | 0.797 | 1.156 | 1.070 | 0.427 |
| Ramachandran plot | | | | |
| Favored (%) | 97.6 | 96.1 | 98.3 | 97.1 |
| Allowed (%) | 2.2 | 3.4 | 1.7 | 2.9 |
| Outliers (%) | 0.2 | 0.6 | 0.0 | 0.0 |
| PDB accession code | 9D2Q | 9D2R | 9D2S | 9D2T |

[a] Values in parentheses refer to the highest resolution shell.

**Table S3. Macromolecular properties of IlvA and IlvA_R from SEC-SAXS**

| Sample | MW[a] (kDa) | MW[b] (kDa) | R*g*[c] (Å) | R*g*[d] (Å) | R*g*[e] (Å) | Res[f] (Å) | Dmax[g] (Å) |
|---|---|---|---|---|---|---|---|
| EcIlvA | 58.2 | 239.6 | 44.76 ± 0.05 | 45.58 ± 0.05 | 45.02 | 54.1 | 156.0 |
| EcIlvA_R | 22.8 | 42.4 | 23.47 ± 0.03 | 23.39 ± 0.03 | 23.15 | 30.6 | 80.0 |
| SaIlvA | 48.0 | 320.9 | 50.95 ± 0.24 | 51.87 ± 0.2 | 51.26 | 67.5 | 171.0 |

[a] Theoretical monomeric molecular weight of the protein.
[b] Molecular weight in solution obtained by Shape&Size (ATSAS) molecular weight analysis.
[c] Radius of gyration ± standard deviation derived from Guinier analysis.
[d] Radius of gyration ± standard deviation derived from GNOM analysis.
[e] Radius of gyration ± standard deviation derived from DENSS analysis.
[f] Fourier Shell Correlation Resolution of the reconstruction from DENSS.
[g] Maximum particle dimension derived from the pair distance distribution function.

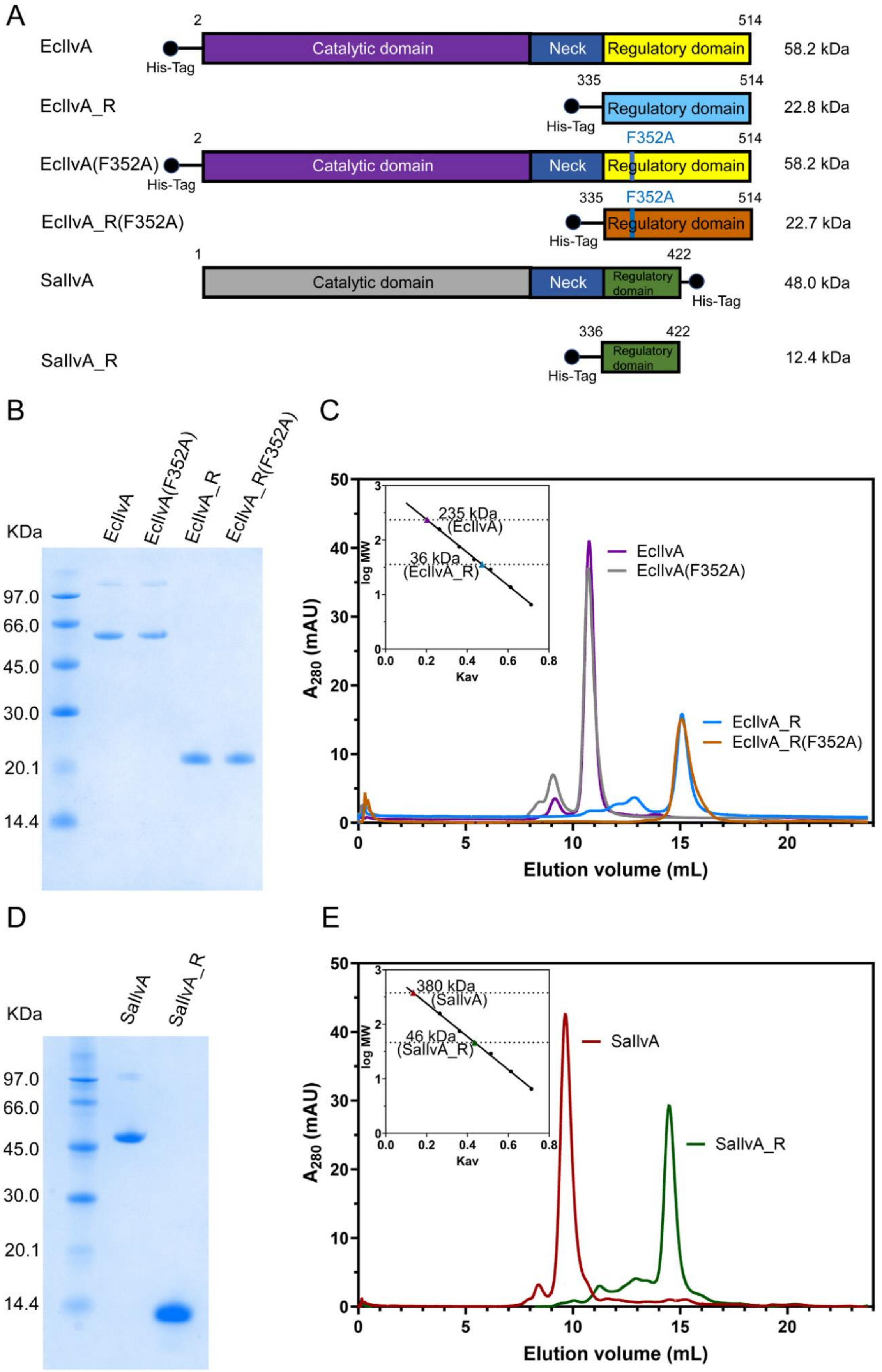
A
EcIlvA
His-Tag
2
514
Catalytic domain
Neck
Regulatory domain
58.2 kDa
EcIlvA_R
335
514
Regulatory domain
22.8 kDa
EcIlvA(F352A)
F352A
58.2 kDa
EcIlvA_R(F352A)
22.7 kDa
SaIlvA
1
422
48.0 kDa
SaIlvA_R
336
12.4 kDa
B
KDa
EcIlvA
EcIlvA(F352A)
EcIlvA_R
EcIlvA_R(F352A)
97.0
66.0
45.0
30.0
20.1
14.4
C
235 kDa (EcIlvA)
36 kDa (EcIlvA_R)
log MW
Kav
A280 (mAU)
Elution volume (mL)
D
SaIlvA
SaIlvA_R
E
380 kDa (SaIlvA)
46 kDa (SaIlvA_R)

**Figure S1. Design and purification of IlvA protein constructs.** *A*, diagrams of the six IlvA protein constructs used in this study. The IlvA protein has an amino terminal catalytic domain connected by a neck to the carboxyl terminal regulatory domain. The presented theoretical molecular weight for the monomer of each construct is calculated by CLC Main Workbench 23.0.2 (QIAGEN). Phe352 is located within the regulatory domain of EcIlvA. *B*, SDS gel electrophoresis showing the purity of the EcIlvA protein constructs. The 120 kDa band is an EcIlvA dimer. *C*, analytical gel filtration chromatography of the four EcIlvA protein constructs using a Superdex 200 10/300 column. *Inset*, the calibration of the gel filtration column with globular protein standards shows EcIlvA (purple) and EcIlvA(F352A) (gray) proteins exist as tetramers, and EcIlvA_R (blue) and EcIlvA_R(F352A) (brown) proteins exist as dimers. *D*, SDS gel electrophoresis showing the purity of the SaIlvA protein constructs. *E*, analytical gel filtration chromatography of the two SaIlvA protein constructs using a Superdex 200 10/300 column. *Inset*, the calibration of the gel filtration column with globular protein standards shows SaIlvA (red) and SaIlvA_R (green) proteins exist as octamer and tetramer, respectively.

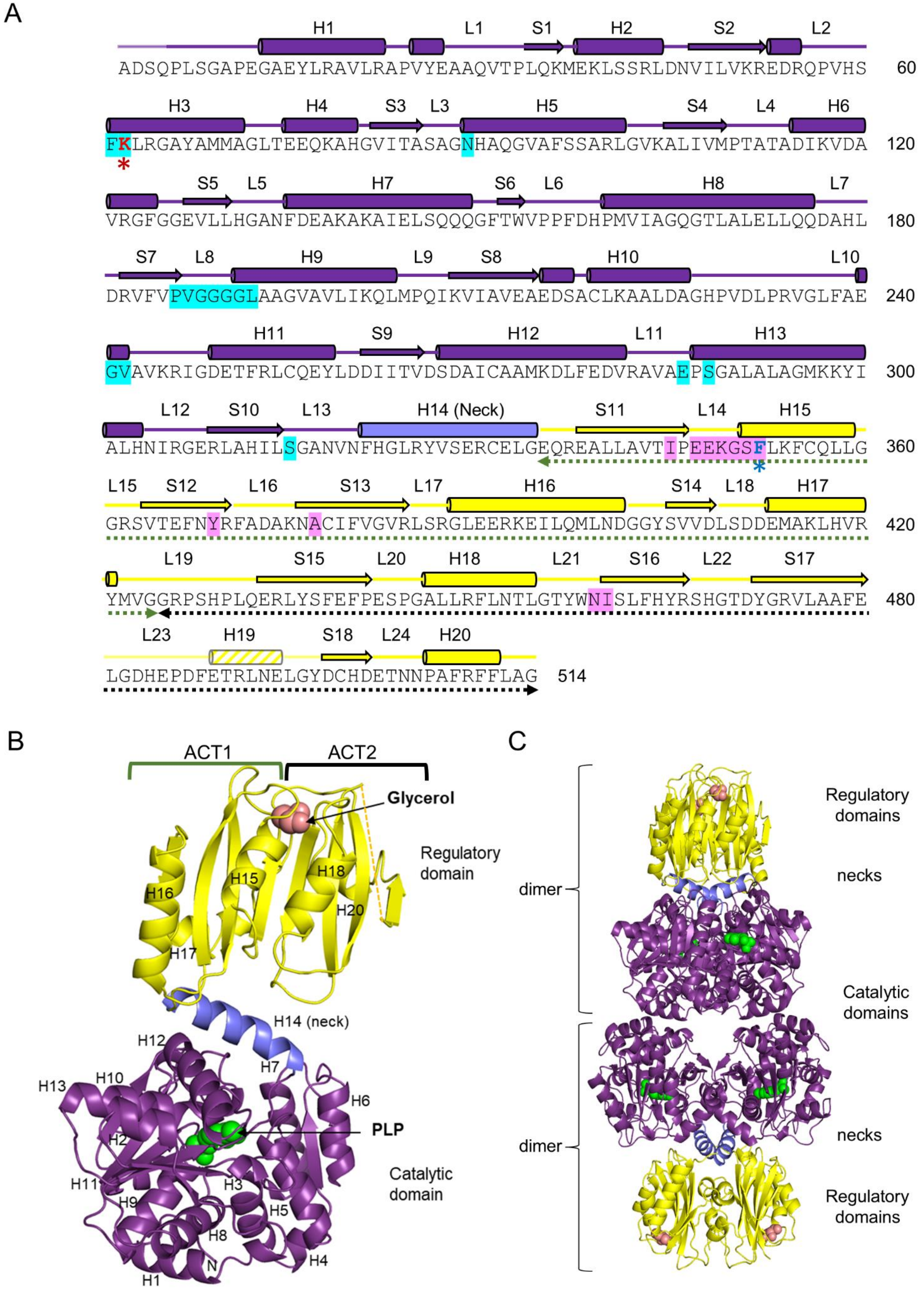
A
H1 L1 S1 H2 S2 L2
ADSQPLSGAPEGAEYLRAVLRAPVYEAAQVTPLQKMEKLSSRLDNVILVKREDRQPVHS 60
H3 H4 S3 L3 H5 S4 L4 H6
FKLRGAYAMMAGLTEEQKAHGVITASAGNHAQGVAFSSARLGVKALIVMPTATADIKVDA 120
S5 L5 H7 S6 L6 H8 L7
VRGFGGEVLLHGANFDEAKAKAIELSQQQGFTWVPPFDHPMVIAGQGTLALELLQQDAHL 180
S7 L8 H9 L9 S8 H10 L10
DRVFVPVGGGGLAAGVAVLIKQLMPQIKVIAVEAEDSACLKAALDAGHPVDLPRVGLFAE 240
H11 S9 H12 L11 H13
GVAVKRIGDETFRLCQEYLDDIITVDSDAICAAMKDLFEDVRAVAEPSGALALAGMKKYI 300
L12 S10 L13 H14 (Neck) S11 L14 H15
ALHNIRGERLAHILSGANVNFHGLRYVSERCELGEQREALLAVTIPEEKGSFLKFCQLLG 360
L15 S12 L16 S13 L17 H16 S14 L18 H17
GRSVTEFNYRFADAKNACIFVGVRLSRGLEERKEILQMLNDGGYSVVDLSDDEMAKLHVR 420
L19 S15 L20 H18 L21 S16 L22 S17
YMVGGRPSHPLQERLYSFEFPESPGALLRFLNTLGTYWNISLFHYRSHGTDYGRVLAAFE 480
L23 H19 S18 L24 H20
LGDHEPDFETRLNELGYDCHDETNNPAFRFFLAG 514
B
ACT1
ACT2
Glycerol
Regulatory domain
H16
H15
H18
H20
H17
H14 (neck)
H12
H7
H13
H10
H6
PLP
H2
Catalytic domain
H11
H3
H9
H5
H8
H4
N
H1
C
Regulatory domains
necks
dimer
Catalytic domains
dimer
necks
Regulatory domains

**Figure S2. Structural overview of EcIlvA.** *A*, the primary structure of EcIlvA with the secondary structural elements labeled. EcIlvA consists of two major domains: the catalytic domain (purple) with PLP bound to Lys62 (*) and the regulatory domain (yellow). Helix H14 (the neck; blue) connects the catalytic and regulatory domains. Two ACT domains in the regulatory domain are shown in green and black dashed arrows. Phe352 (*) is located in the regulatory domain. The residues highlighted in cyan are involved with PLP binding and the residues highlighted in magenta are involved with isoleucine binding. Residues 482-496 are disordered (dotted line). *B*, a labeled secondary structure representation of EcIlvA colored as in Panel A. PLP is shown in green, and glycerol is shown in salmon. *C*, the crystal structure of the EcIlvA tetramer.

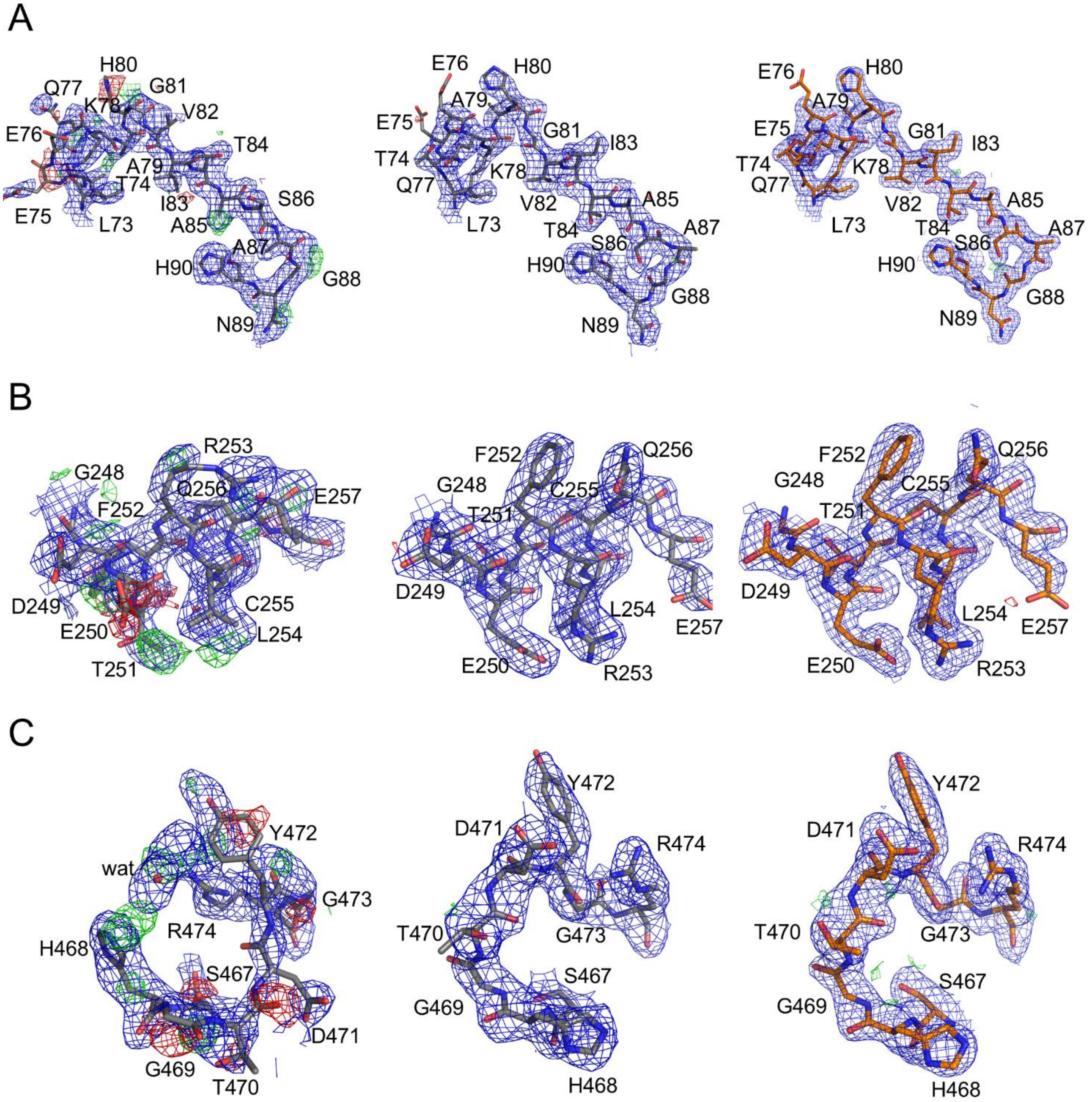


**Figure S3. Improved model in three regions of EcilvA strcuture.** Electron density maps with model structures of regions 1 (*A*), 2 (*B*), and 3 (*C*) from Figure 2*A*. Left panels show the published EcIlvA model with 2Fo-Fc and Fo-Fc electron density maps downloaded from PDB (PDB ID 1TDJ). Middle panels show the re-fitted and refined model with 2Fo-Fc and Fo-Fc electron density maps. The positive and negative electron densities have disappeared in the Fo-Fc electron density maps. Right panel shows the higher resolution structure with 2Fo-Fc and Fo-Fc electron density maps (PDB ID 9D2Q). The 2Fo-Fc electron density maps (blue mesh) are contoured at 1.0σ, and the Fo-Fc electron density maps are countoured at 3.0σ (green mesh) and **-**3.0σ (red mesh).

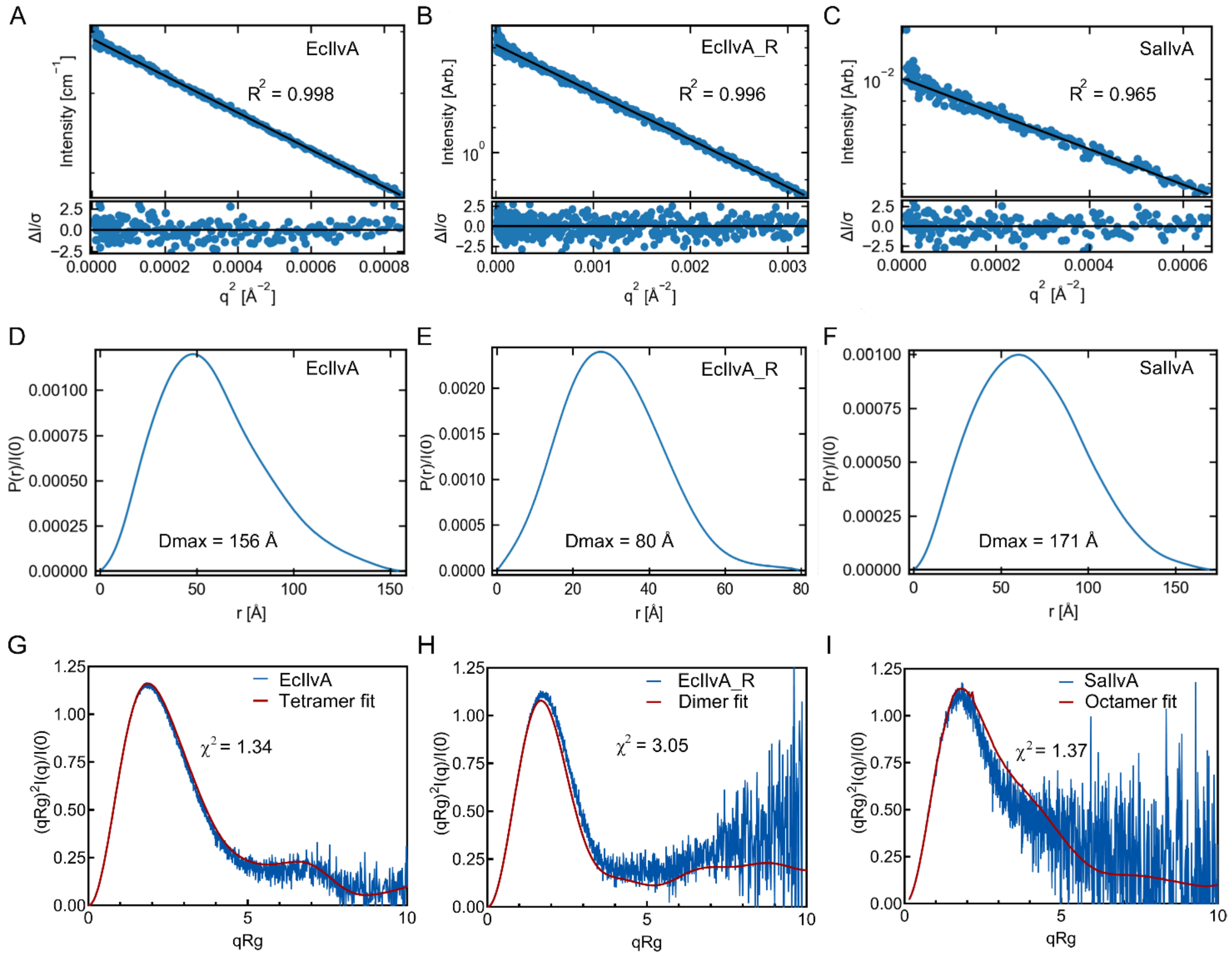


**Figure S4. Analysis of the IlvA solution dynamics by SEC-SAXS.** *A-C*, Guinier fit (top) and fit residuals (bottom) of EcIlvA, EcIlvA_R and SaIlvA. *D-F*, P(r) function of EcIlvA, EcIlvA_R and SaIlvA, normalized by I(0). *G*, the dimensionless Kratky plots of EcIlvA SAXS data (blue) compared to the theoretical Kratky plot (red) calculated from a model of the IlvA tetramer (PDB ID 9D2Q) fit into the SAXS volume. *H*, the dimensionless Kratky plots of EcIlvA_R SAXS data (blue) compared to the theoretical Kratky plot (red) calculated from a model of the EcIlvA_R dimer (PDB ID 9D2R) fit into the SAXS volume. *I*, the dimensionless Kratky plots of SaIlvA SAXS data (blue) compared to the theoretical Kratky plot (red) calculated from a model of the SaIlvA octamer fit into the SAXS volume. The octamer is assembled with two tetramers. The model of the SaIlvA tetramer is generated using an AlphaFold model of the SaIlvA dimer and crystal structure of SaIlvA_R (PDB ID 9D2T). The $\chi^2$ values reflecting the fit between the theoretical model and the SAXS data are shown in each panel.

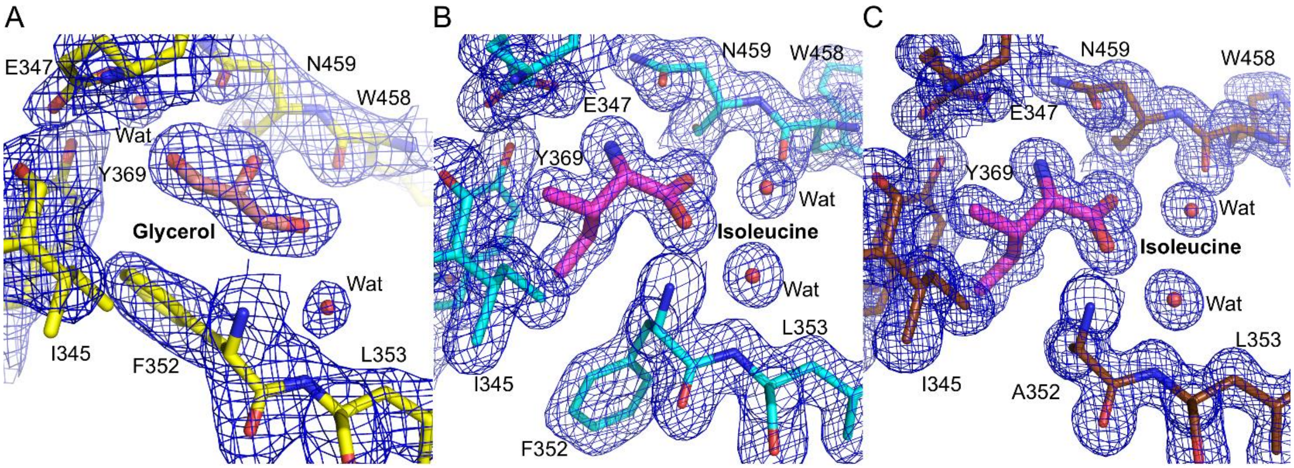


**Figure S5. Electron density maps.** *A*, the 2Fo-Fc electron density map of bound glycerol, two associated water molecules, and the key residues that interact with glycerol in the EcIlvA structure (PDB ID: 9D2Q). *B*, the 2Fo-Fc electron density map of bound isoleucine, two water molecules, and the key residues that interact with isoleucine in the EcIlvA_R•Ile complex structure (PDB ID: 9D2R). *C*, the 2Fo-Fc electron density map of bound isoleucine, two water molecules, and the key residues that interact with isoleucine in the EcIlvA_R(F352A)•Ile complex structure (PDB ID: 9D2S). The 2Fo-Fc electron density maps (blue mesh) are contoured at 1.0σ.

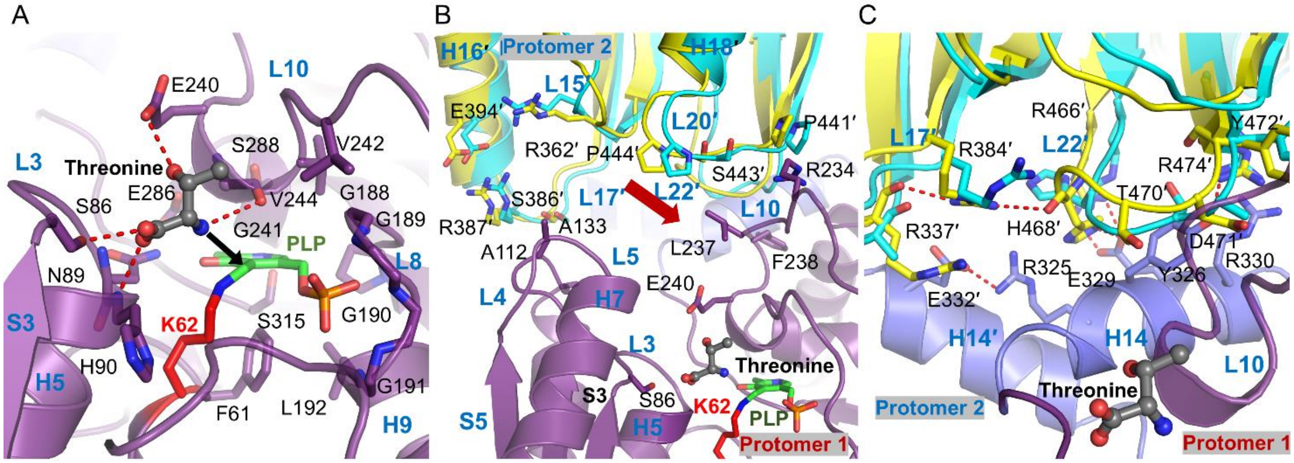


**Figure S6. Threonine binding at the catalytic site.** *A*, threonine docked into the EcIlvA active site (PDB ID: 9D2Q). Threonine lies between loops L3 and L10 of the catalytic domain (purple). The molecular docking solution was calculated using the Molecular Operating Environment software (2018.01, Chemical Computing Group). The interaction between the amine group of threonine and the aldehyde carbon atom of PLP is shown as a black arrow. *B*, change in loop interactions between the catalytic and regulatory domain by the binding of isoleucine. The conformational change of the loop L20 is shown as a red arrow. *C*, change in interactions between the necks and regulatory domain by the binding of isoleucine. The catalytic domain of protomer 1 and regulatory domain of protomer 2 of EcIlvA are shown in purple and yellow, respectively. The overlaid EcIlvA_R•Ile is shown in cyan. Residues on the opposite protomer (promoter 2) are indicated with a prime. The docked threonine molecule is represented by ball-and-sticks with gray carbon atoms, blue nitrogen atom, and red oxygen atoms. PLP and the active-site lysine (K62) are shown as sticks.

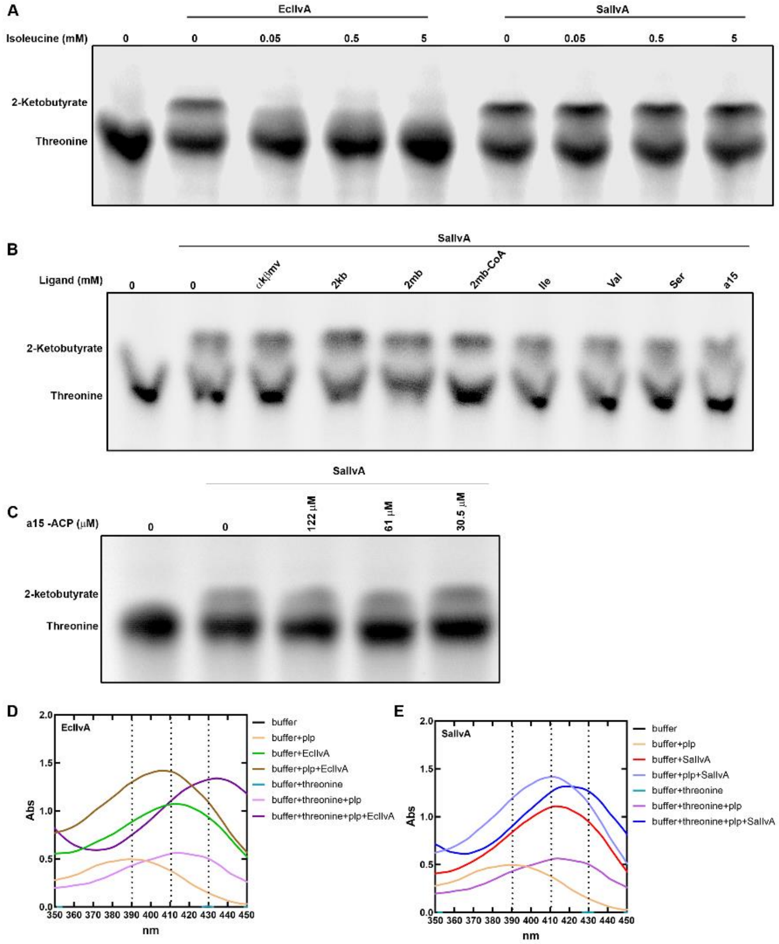


**Figure S7. EcIlvA and SaIlvA assay with threonine.** *A*, TLC showing inhibition of 14[C] 2-ketobutyrate formation from 14[C] threonine by EcIlvA or SaIlvA by isoleucine. *B*, TLC showing 14[C] 2-ketobutyrate formation from 14[C] threonine by SaIlvA in the presence of intermediates in the branched chain amino acid pathway. αkβmv (alpha-keto-beta-methylvalerate) (IlvE substrate) - 5 mM, 2kb (2-ketobutyrate) (IlvA product) – 5 mM, 2mb (2-methylbutyrate) – 5 mM, 2mb-CoA (2-metylbutyryl-CoA) – 4 mM, Ile (Isoleucine) – 5 mM, Val (Valine) – 5 mM, Ser (Serine) – 5mM, and a15 (anteiso-15 fatty acid) - 150 µM. *C*, TLC showing 14[C] 2-ketobutyrate formation from 14[C] threonine by SaIlvA in the presence of anteiso15-ACP (a15-ACP). *D*, Spectra of EcIlvA in the presence of 0.1 mM plp or 0.1 mM plp + 30 mM threonine. *E*, Spectra of SaIlvA in the presence of 0.1 mM plp or 0.1 mM plp + 30 mM threonine.

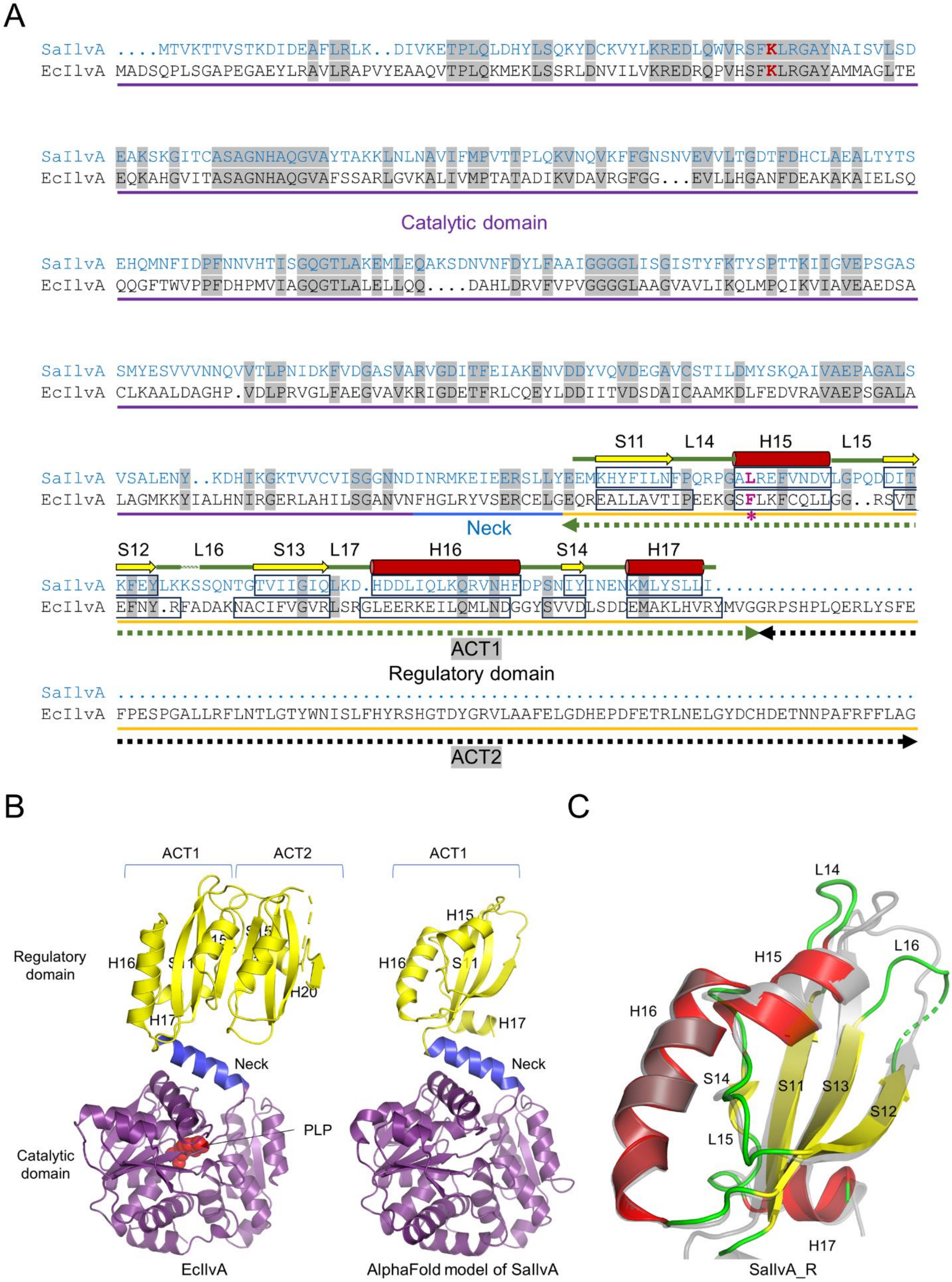
A
SaIlvA ....MTVKTTVSTKDIDEAFLRLK..DIVKETPLQLDHYLSQKYDCKVYLKREDLQWVRSFKLRGAYNAISVLSD
EcIlvA MADSQPLSGAPEGAEYLRAVLRAPVYEAAQVTPLQKMEKLSSRLDNVILVKREDRQPVHSFKLRGAYAMMAGLTE
SaIlvA EAKSKGITCASAGNHAQGVAYTAKKLNLNAVIFMPVTTPLQKVNQVKFFGNSNVEVVLTGDTFDHCLAEALTYTS
EcIlvA EQKAHGVITASAGNHAQGVAFSSARLGVKALIVMPTATADIKVDAVRGFGG...EVLLHGANFDEAKAKAIELSQ
Catalytic domain
SaIlvA EHQMNFIDPFNNVHTISGQGTLAKEMLEQAKSDNVNFDYLFAAIGGGGLISGISTYFKTYSPTTKIIGVEPSGAS
EcIlvA QQGFTWVPPFDHPMVIAGQGTLALELLQQ....DAHLDRVFVPVGGGGLAAGVAVLIKQLMPQIKVIAVEAEDSA
SaIlvA SMYESVVVNNQVVTLPNIDKFVDGASVARVGDITFEIAKENVDDYVQVDEGAVCSTILDMYSKQAIVAEPAGALS
EcIlvA CLKAALDAGHP.VDLPRVGLFAEGVAVKRIGDETFRLCQEYLDDIITVDSDAICAAMKDLFEDVRAVAEPSGALA
S11 L14 H15 L15
SaIlvA VSALENY..KDHIKGKTVVCVISGGNNDINRMKEIEERSLLYEEMKHYFILNFPQRPGALREFVNDVLGPQDDIT
EcIlvA LAGMKKYIALHNIRGERLAHILSGANVNFHGLRYVSERCELGEQREALLAVTIPEEKGSFLKFCQLLGG..RSVT
Neck
*
S12 L16 S13 L17 H16 S14 H17
SaIlvA KFEYLKKSSQNTGTVIIGIQLKD.HDDLIQLKQRVNHFDPSNIYINENKMLYSLLI...................
EcIlvA EFNY.RFADAKNACIFVGVRLSRGLEERKEILQMLNDGGYSVVDLSDDEMAKLHVRYMVGGRPSHPLQERLYSFE
ACT1
Regulatory domain
SaIlvA ..........................................................................
EcIlvA FPESPGALLRFLNTLGTYWNISLFHYRSHGTDYGRVLAAFELGDHEPDFETRLNELGYDCHDETNNPAFRFFLAG
ACT2
B
ACT1
ACT2
ACT1
Regulatory domain
H16
H17
H20
H15
H16
H17
Neck
Neck
PLP
Catalytic domain
EcIlvA
AlphaFold model of SaIlvA
C
L14
H15
L16
H16
S14
S11
S13
S12
L15
H17
SaIlvA_R

**Figure S8. Structural overview of SaIlvA_R.** *A*, the sequence alignment of SaIlvA and EcIlvA was performed with UCSF Chimera. IlvA consists of two major domains: the catalytic domain (purple underline) with PLP bound to a lysine (red), the regulatory domain (yellow) and the neck (blue) that connects the catalytic and regulatory domains. Light grey regions show residues conserved in both sequences. The secondary structural elements of SaIlvA_R (PDB ID: 9D2T) are depicted above the sequence alignment according to EcIlvA labels. Unlike EcIlvA_R, which is composed of two ACT domains (ACT1 and ACT2), SaIlvA_R is composed of only one ACT domain (ACT1). ACT1 and ACT2 domains are shown with green and black dashed underline, respectively. Phe352 of EcIlvA and its corresponding residue Leu352 of SaIlvA are indicated in magenta with an asterisk. *B*, AlphaFold model structure of SaIlvA (AF-Q2FF63) compared with EcIlvA (PDB ID: 9D2Q). The regulatory domain of EcIlvA has two ACT like domains (left), while SaIlvA contains only one ACT like domain in regulatory domain (right). The RMSD between two regulatory domains is 3.5 Å on 83 Cα atoms. *C*, the SaIlvA_R monomer (PDB ID: 9D2T) is superimposed on the AlphaFold model of SaIlvA (gray) and colored as in Panel A. The secondary structures, except for loops L14 and L16, superimpose well.

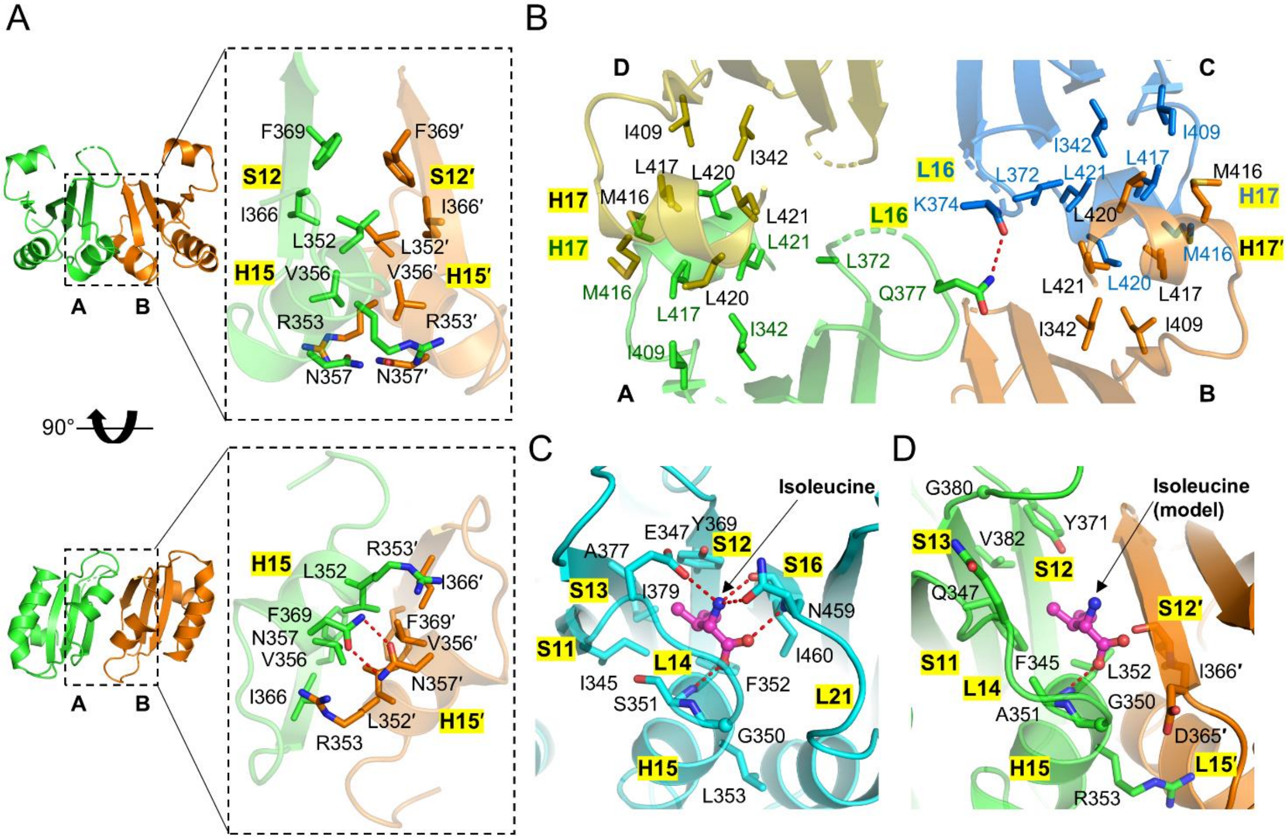


**Figure S9. The dimeric structure of SaIlvA_R.** *A*, dimeric SaIlvA_R corresponds to the monomer of EcIlvA_R and was made by using protomers A and B from the crystal structure of SaIlvA_R. The protomers A and B are related to two-fold symmetry and are shown with green and orange ribbon, respectively. The view is rotated by 90° (below). Close-up views of the dimer interface are shown with key interaction residues (right). *B*, close-up view of the dimer-dimer interface of the SaIlvA_R tetramer. The major clusters of hydrophobic residues in the dimer-dimer interface are shown as stick models. The protomers within the tetramer of SaIlvA_R are shown in green (protomer A), orange (protomer B), blue (protomer C), and yellow (protomer D). *C-D,* superimposed structures on helix H15 of EcIlvA_R(C) and SaIlvA_R(D). Sheet S16 of EcIlvA_R (C) corresponds to S12′ of SaIlvA_R (D). Protomers A and B of SaIlvA are shown in green and orange, respectively. The features of protomer B are indicated with a prime.

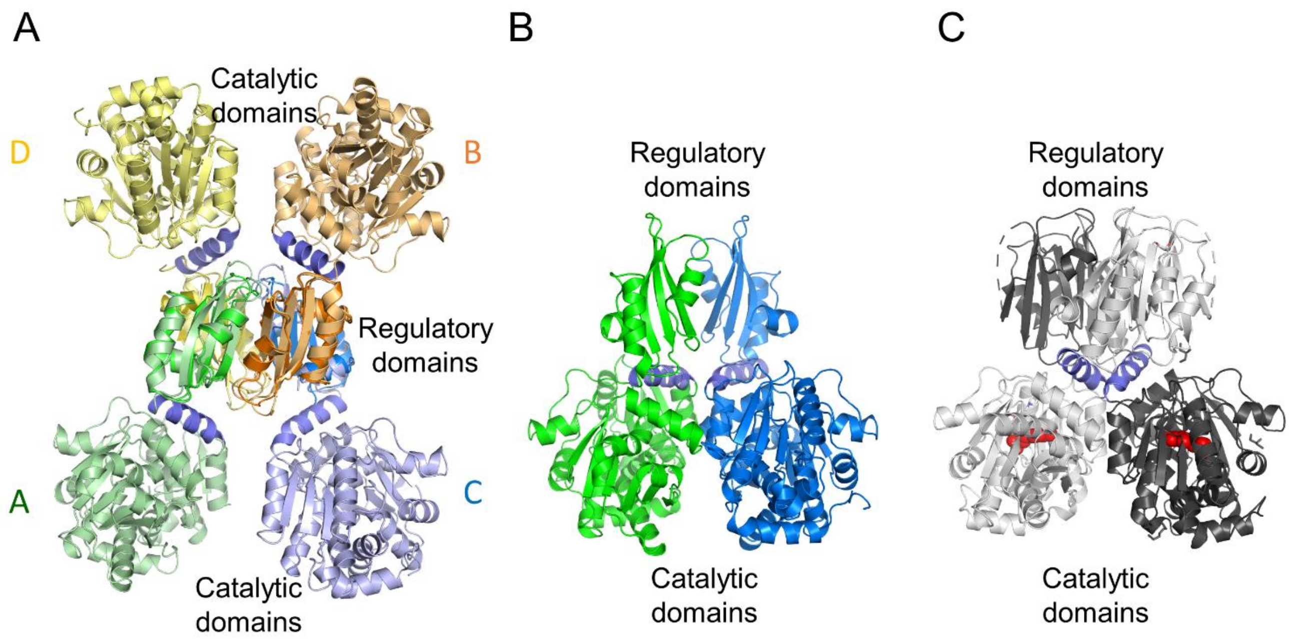


**Figure S10. AlphaFold model of SaIlvA.** *A*, overlaid SaIlvA AlphaFold models on tetrameric SaIlvA_R are shown in green (protomer A), orange (protomer B), blue (protomer C), and yellow (protomer D). There is no close contact between the catalytic domains. *B*, dimer of SaIlvA generated using AlphaFold. *C*, crystal structure of EcIlvA dimer.

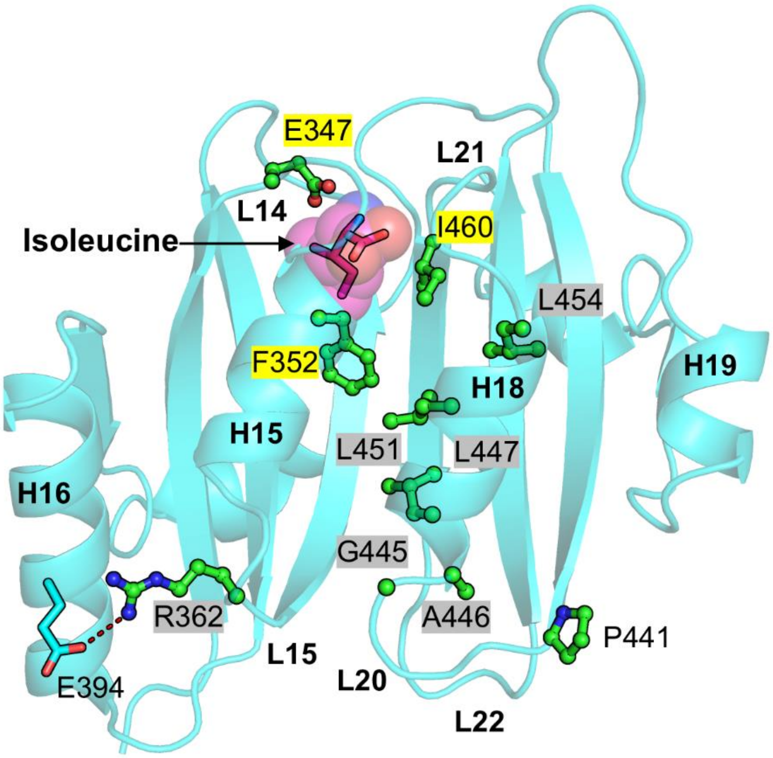


**Figure S11. Structural location of mutations reported to render EcIlvA refractory to isoleucine inhibition.** The mutated residues of EcIlvA are mapped onto the EcIlvA_R•Ile crystal structure (PDB ID: 9D2R). The residues that directly interact with isoleucine are highlighted in yellow, and the residues that are proposed to relay the isoleucine binding signal to loops L15 and L20 are highlighted in gray.
